# Deep learning representations of human Immune Health for precision immunology

**DOI:** 10.64898/2026.07.15.724605

**Authors:** Matthew E. Lee, Jaesik Kim, Matei Ionita, Michelle L. McKeague, Jonghyun Lee, Yonghyun Nam, Chang-Uk Jeong, Akira Nair, Edmore T. Moyo, Irene Khavin, Kevin Wang, Shwetank, Divij Mathew, Victoria Fang, Benjamin A. Fensterheim, Ajinkya Pattekar, Zahabia Rangwala, Amit Bar-Or, Benjamin A. Abramoff, Rennie L. Rhee, Stephen Schuster, Alexander Huang, Nuala Meyer, Maayan Levy, Alfred Garfall, Vijay Bhoj, Mary Kaminski, Ali Naji, Enjun Yang, Christopher Cabanski, John Connolly, Leonardo Guercio, Joost Wagenaar, Amy E. Baxter, Damian Maseda, Sokratis A. Apostolidis, Mark M. Painter, Robert H. Vonderheide, Allison R. Greenplate, Dokyoon Kim, E. John Wherry

## Abstract

The human immune system is composed of ∼30-50 distinct cell types, each of which can exist in different states of activation or differentiation. Indeed, the mammalian immune system has evolved to sense and respond to infections, cancers, injuries, and changes in tissue or host homeostasis (1). Moreover, an increasingly large fraction of approved drugs target the immune system directly, and/or cause immune changes (2–4). A key feature of the immune system is to store some of this information, for example as innate or adaptive immune memory (5). In addition, rewiring of immune network architecture induced by disease, environmental exposures, drug treatments, and/or chronological age allows the immune system to store information in the pattern of connections and activity across populations of immune cells. This ensemble information storage, in addition to changes to individual cells, functions as a major way the immune system encodes aspects of immune history and future potential. Genetic information can identify inherited risk alleles, but cannot capture the continual remodeling of the immune system shaped by exposures, infection, inflammation, therapy, and aging (6, 7). To define and use such ensemble immunotypes, we developed a self-supervised deep learning framework that transforms high-dimensional immune profiles into representations of immune health. MAESTRO (MAsked Encoding Set TRansformer with self-distillatiOn) encodes a set of cells from an individual into an embedding that captures immune cell population-level organization. Pretrained on 1,792 peripheral blood samples comprising over 418 million immune cells across 13 clinical diagnoses, MAESTRO learns immune fingerprints that are stable within individuals yet diverse across populations, states of health, disease, and treatment, providing a quantitative basis for comparing immune states across individuals and over time. These fingerprints capture immune architecture beyond coarse cell type proportions, enabling patient-efficient clinical prediction using simple task specific models. MAESTRO model embeddings retain a temporal dimension of immune history and potential, reflecting signatures of past exposures and baseline features that predict future immune responses. Finally, we demonstrate a translational precision immunotherapy application by testing this approach in metastatic Pancreatic Ductal Adenocarcinoma (PDAC), where pretreatment immune landscape circuitry maps enable patient stratification and therapeutic response prediction. Overall, we developed a large, attention-based model that captures deep network architecture of immune states through self-supervised representations of immune cytometry data as a reusable foundation for precision immunology, converting immune complexity into clinically actionable embeddings for diagnosis, monitoring, and therapy selection.

## Main

The mammalian immune system evolved to sense and respond to a diverse set of perturbations of host tissue and systemic homeostasis (1). Immune responses during organismal development, colonization with microbes, injury, infection, cancers, environmental exposures, and autoreactivity, involve dozens of immune cell types, each endowed with complex and diverse sets of activities, functions, and roles. The activity (or products) of immune cells can sometimes point to disease mechanisms and/or drug targets (8–10). However, the complexity of immune states and future responsiveness is also encoded in the interactions *between* cells much like cognitive memories are stored by ensembles of neurons in addition to changes in individual cells (11–13). Changes to the system architecture of immune networks may, therefore, be an important non-genetic determinant of future immune responses based on previous immune exposures. Indeed, while previous immune exposures can be stored as immune memories for individual cell types (innate or adaptive), the rewiring of the overall immune system landscape, while appreciated for selected events (e.g. CMV infection (7)), has been less-well interrogated for predicting future immune responses.

Precision medicine, one of the primary objectives of modern biomedical research, requires quantitative representations of patient-specific biology to enable patient-specific treatments. Genomic sequencing has advanced these approaches, particularly in cancer where tumor mutations can be drugged and in some rare diseases where germline mutations can be corrected (14–16). However, static DNA sequence alone cannot efficiently capture changes to host biology caused by environmental exposures, infections, inflammation, and aging (6, 7). The immune system, however, continually integrates information about immune responses and immune perturbations across lifespan, forming and maintaining immune network changes, establishing immune memory, and adapting to new encounters (17, 18). The immune system is implicated in many if not most diseases (19–21) suggesting that defining patient-specific immune architecture patterns may help define disease relevant immune fingerprints and enable precision therapeutics based on patient immunotype.

In healthy individuals, immune cell networks have stable patterns, with greater variation between people than within the same person over time (22, 23). This stability supports the idea that human immune systems are built from the same components yet become uniquely structured in each individual, consistent with the notion of an immune “fingerprint” that is shaped by both genetics and immune history (7, 22–24). Whether variation in these immune fingerprints can be leveraged to better understand immune health, predict disease risk, and/or inform treatment decisions remains an open question (25). Capitalizing on the potential of immune fingerprints to predict disease trajectories or therapeutic responses requires measuring immune complexity across individuals at scale.

Understanding any biological system requires interrogating both the individual components of the system and the relationships that connect these components into network-like structures. For example, recent deep learning approaches model the single cell as a system, leveraging the whole transcriptome resolution of RNA sequencing as units of the system to represent intracellular complexity (26–35). To encode information about how individual cells work, scaling for transcriptomic depth will enable deep learning of cell behavior. For the immune system, although the intracellular circuitry of individual cells in isolation is important, the function and behavior of the system depends on the relationships *between* immune cell populations and the dynamic reorganization of these network interactions through health and disease (8, 11–13, 36). To encode information about individual human immune systems, scaling measurements to large numbers of cells and relationships between them should empower deep learning about the behavior of entire immune systems from individual subjects and patients. In other words, prioritizing number of cells per sample timepoint rather than depth of information for any individual cell enables learning system behavior compared to individual cell behavior. Flow and mass cytometry make it possible to collect data on hundreds of thousands to millions of immune cells per sample, capturing the compositional complexity and population structures that define individual-level immune architecture (37). Translating these raw measurements into quantitative representations faces fundamental computational challenges, including the need to scale methods to transform cytometry and/or immune cell data into quantitative immune representations for precision immune health.

One approach to address some of these computational needs is Deep Representation Learning (DRL). DRL has emerged as a powerful framework for structuring complex biological data (38–41). Unlike supervised learning that requires annotated training data or traditional unsupervised methods that identify structure without explicit annotation-guided training objectives, self-supervised DRL uses the data itself as both input and training signal, enabling models to learn meaningful biological patterns from large unannotated datasets. While previous DRL approaches for cytometry and single-cell data have focused on characterizing individual cells, quantitatively representing an individual immune system requires encoding sets of cell populations as an embedding (26–35, 42). The shift from cell-level to sample-level encoding is necessary because immune function emerges as the ensemble or the collective behavior of multiple cell populations, not from individual immune cell types in isolation. Such learned embeddings of the collective system of multiple immune cell types operating in concert should enable quantitative comparisons of immune states across individuals and timepoints, providing the framework to translate unique immune structure variation into clinical insights. Such an approach could transform conventional immunology into translationally actionable immune health capable of informing treatment selection, predicting therapeutic response, and tracking health and disease changes in real-time.

Here, we developed an attention-based Immune Health model and scalable framework for quantitative immune system representation using large-scale cytometric immune profiling data that addresses these challenges. We developed a two-stage approach that decouples learning immune fingerprints from task-specific predictions. We trained a self-supervised, representation learning model on a large corpus of unannotated cytometry data to learn fundamental patterns of immune structure. This model processes entire blood-based cytometry samples into embeddings, preserving all cellular and overall compositional information. The learned embeddings capture biologically meaningful structure, including disease-specific patterns that associate with disease diagnosis and stable individual-specific immune signatures that persist over time. These embeddings encode information spanning temporal scales including signatures of past exposures (e.g. in utero HIV exposure) and predictions of future immune responses (e.g. vaccine response). We also translated this approach to precision immunotherapy, demonstrating prospective prediction of treatment response in pancreatic cancer patients. This scalable framework transforms cytometry data into clinically actionable immune fingerprints, providing the quantitative foundation for immune-based precision medicine.

### A scalable framework for dynamic precision medicine using Immune Health

Unlike germline encoded genes, the human immune system is continuously remodeled in response to environmental exposures, microbes and infections, injuries, and tissue and organismal aging. As a result, immune profiles that reflect genetics and non-genetic effects of previous exposures are well-suited for precision medicine applications (**Fig. 1a**). We assembled a high-dimensional cytometry dataset of blood-based immune cells comprising 1,792 samples across multiple clinical cohorts, each with associated clinical metadata (**Fig. 1b**). We employed mass cytometry (CyTOF) (37) to capture data for large numbers of individual cells, and the relationships between these cells in each sample (**Fig. 1b**). The advantage of cytometry is the capacity to collect data from hundreds of thousands of cells per sample, prioritizing more features per sample (cells) over smaller numbers of cells with more features per cell (e.g. scRNA-seq).

**Figure 1:**
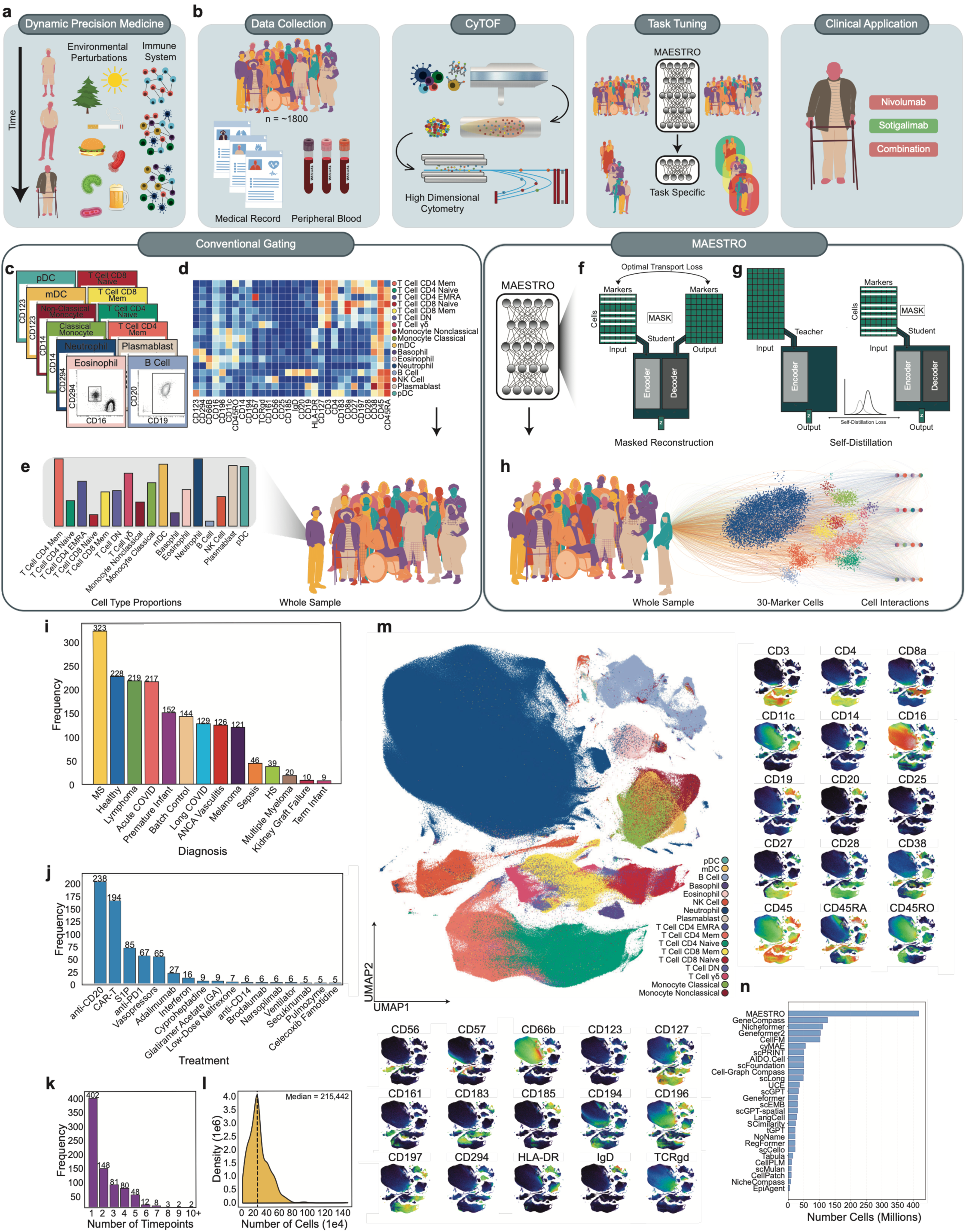
A scalable framework for dynamic precision medicine from cytometry. **a.** The immune system continuously remodels across the lifespan, incorporating various perturbations. **b.** Data Collection: cohort assembly from peripheral blood samples linked to clinical metadata (n = 1,792). CyTOF: workflow for high-dimensional single-cell immune profiling. Task Tuning: two-stage framework: self-supervised pretraining of MAESTRO on unannotated cytometry data, followed by task- specific models trained on frozen embeddings. Clinical Application: metastatic PDAC, where baseline immune fingerprints inform treatment selection. **c.** Conventional hierarchical gating strategy (17 populations; **Supplementary Fig. 1**). **d.** Median marker expression per gated population. **e.** Conventional sample representation as a cell-type proportion vector. **f.** Masked reconstruction objective: cells are masked and reconstructed from remaining cellular context via optimal transport loss. **g.** Self-distillation objective: a Teacher encodes the full cell set while a Student encodes masked subsampled views, with representations aligned via self-distillation loss. **h.** MAESTRO encodes whole samples from 30-marker single-cell measurements into compact embeddings preserving population- level structure and cell-interaction information. **i.** Diagnosis distribution across the pretraining corpus. **j.** Treatment exposure distribution. **k.** Number of longitudinal timepoints per individual. **l.** Cells per sample distribution (median = 215,442). **m.** UMAP of 5 million cells colored by immune population with marker intensity overlays for all 30 markers. **n.** Training corpus size compared to other single-cell foundation models.

The high dimensionality of such immune profiling data combined with the challenges and costs of recruiting large numbers of human subjects means that clinical cohorts are often too small to train robust predictive models without overfitting. We addressed this issue with a two-stage framework that decouples general immune fingerprint learning from task-specific immunotype identification (**Fig. 1b**). First, MAESTRO is trained on a large, unannotated cytometry dataset generated from multiple cohorts through self-supervised learning. This approach transforms raw cytometry measurements into structured embeddings that are fixed-length numerical vectors, capturing patterns of immune organization. Second, smaller task-specific models are trained using these embeddings as input to identify clinically relevant patterns in smaller clinical cohorts. This approach leverages the notion that immune systems share common organizational principles despite variation between individuals. MAESTRO learns these shared principles from large-scale data, and task-specific models resolve meaningful variation in smaller cohorts. New patients are processed through both models sequentially, where raw cytometry data is first encoded into embeddings by MAESTRO, then mapped to immunotypes for applications such as personalized treatment selection (**Fig. 1b**). This framework transforms individual immune complexity into a robust foundation for precision medicine.

Immune profiles are conventionally constructed by manually annotating cytometry data, where cells are sequentially partitioned into discrete cell-populations based on marker expression patterns (**Fig. 1c, Supplementary Figure 1**). For example, **Fig 1d** shows marker expression for different immune cell types. Immune profiles from such manually annotated immune data are represented as a vector of immune cell-population frequencies (**Fig. 1e**). However, this approach reduces continuous marker distributions to categorical assignments, ignores relationships between cells and markers, and scales poorly with increasing panel complexity and number of samples. These limitations suggest alternative approaches might not only scale more easily but also capture additional immune information.

MAESTRO addresses some of these challenges and learns immune fingerprints through two complementary self-supervised training strategies, without the need to identify cell populations. In masked reconstruction, MAESTRO strategically hides cells from a sample and then learns to predict marker expression patterns for the missing cells based on the remaining cells (**Fig. 1f**). Because all information must pass through a compressed embedding before reconstruction, the model can only recover masked cells if that embedding captures the relationships between cell populations in the sample. Masked reconstruction therefore forces the embedding to encode immune compositional structure and inter-population dependencies. We then used a self-distillation approach. In self-distillation, two neural networks termed “Student” and “Teacher” process the same immune profile differently: The Teacher model observes all cells to produce a reference embedding, while the Student model processes smaller subsets with masked cells and performs the computationally intensive learning. Training aligns these two representations, forcing the Student to capture stable immune structure from partial cellular information rather than relying on any particular subset of cells. This strategy reinforces robust feature learning while enabling efficient scaling to millions of cells per sample (**Fig. 1g**). Together, these features force MAESTRO to learn robust embeddings of immune organization from large-scale unannotated data.

MAESTRO encodes the learned cellular information into a compact, structured representation of the immune profile of each sample. Rather than comparing every cell to every other cell, MAESTRO learns a set of immune prototypes, and characterizes each cell by its similarity to these learned reference points (Cell Prototype Encoder, **Supplementary Figure 2**). These prototypes capture different sets of immune cell patterns and relationships between cells that define the overall immune landscape distinguishing individual health and disease states. MAESTRO then condenses this information into a single vector by aggregating all processed cells based on their learned importance (Pooling Encoder, **Supplementary Figure 2**), preserving the full distribution of cellular measurements and relationships among cell-populations within a compact representation (**Fig. 1h**).

The goal of MAESTRO was to build an attention-based model capable of resolving individual states of immune health and disease and predict trajectories, including after therapeutic intervention. Training a robust model requires data that is diverse, longitudinal, and large scale. To this end, we assembled a training corpus of standardized high dimensional cytometry data from 1,792 peripheral blood samples comprising over 418 million cells across 13 clinical diagnoses plus batch controls. This dataset captured immune perturbations spanning autoimmunity, infection, malignancy, and transplantation (**Fig. 1b, 1i**).

Diversity is critical to drive learning in models (43). Among the healthy and disease states interrogated, multiple sclerosis (MS) (n=323), healthy donors (n=228), lymphoma (n=219), and (hospitalized) acute COVID-19 (n=217) were the largest cohorts, while smaller cohorts including sepsis, hidradenitis suppurativa (HS), and kidney graft failure provided additional immune diversity. Treatment annotations documented 17 therapeutic categories across these cohorts (**Fig. 1j**), with immunomodulatory agents predominating including anti-CD20 therapies (n=203), CAR T cells (n=194), S1P receptor modulators (n=85), and checkpoint inhibitors including anti-PD1 (n=67). The dataset had balanced sex representation (966 male, 826 female) (**Supplementary Fig. 3a**) and spanned the human lifespan from premature infants to elderly adults (median age 42.8 years), with a bimodal age distribution reflecting the infant and adult cohorts (**Supplementary Fig. 3b**). This heterogeneity exposes MAESTRO to a broad spectrum of immune states, from acute inflammatory responses to chronic immune dysregulation.

In addition to diversity of immune profiles, longitudinal sampling is particularly informative for the immune system and enables encoding of temporal dynamics in learned representations. Among the 1,792 samples, 384 subjects contributed multiple timepoints, with 148 individuals sampled at two timepoints and a smaller number of individuals sampled at 3 or more time points (**Fig. 1k**). Repeated sampling from the same individual exposes MAESTRO to both the stable features that define a patient- or subject-specific immune fingerprint, as well as the dynamic changes that may occur over time. This temporal structure enables MAESTRO to learn representations that distinguish durable individual-level immune architecture from transient perturbation-driven changes.

The use of cytometry enabled acquisition of hundreds of thousands to millions of cells per sample. Each sample contained a median of 215,442 cells, with a distribution spanning tens of thousands to over one million cells per sample (**Fig. 1l**). Visualized through a random subsample of 5 million cells projected via UMAP (**Fig. 1m**), cells organized into clusters corresponding to canonical immune populations. This corpus of immune data exceeds the training corpora of existing single-cell foundation models by approximately fourfold or more (**Fig. 1n**), enabling MAESTRO to overcome the data sparsity constraints that have limited the application of deep learning to patient-level immune profiling.

### Self-supervised learning captures meaningful immune structure

To evaluate whether MAESTRO has learned to embed accurate immune structure, we tested the ability to predict masked cell populations from the remaining unmasked cells across diverse immune states. We visualized this process across samples including healthy controls, acute infection, premature infants, malignancy, and treatment-induced perturbations (**Fig. 2a**). Because immune composition varies across individuals and clinical states, we wanted a model that would not need to rely on a single template but was able to predict missing populations from diverse immune contexts. Thus, MAESTRO used a cell population-level masking strategy that hides phenotypically similar cell populations rather than random cells, forcing the model to infer missing populations from immunologically distinct cells. Indeed, for healthy donors where there were typical distributions of major lineages the model recovers masked T cell and myeloid populations with high fidelity. MAESTRO also recovered immune profiles skewed by biology, disease, or treatment, effectively capturing the developmentally distinct immune architecture of premature infants, the distinct immune landscape in B cell lymphoma patients with peripheral blasts, and the altered immune populations in patients treated with S1P receptor modulator where T cells are sequestered in the lymph node and depleted from peripheral blood (**Fig. 2a**). Across these diverse immunological contexts, MAESTRO demonstrated robust predictive capacity, indicating that self-supervised training captured generalizable principles of immune cell population organization.

**Figure 2:**
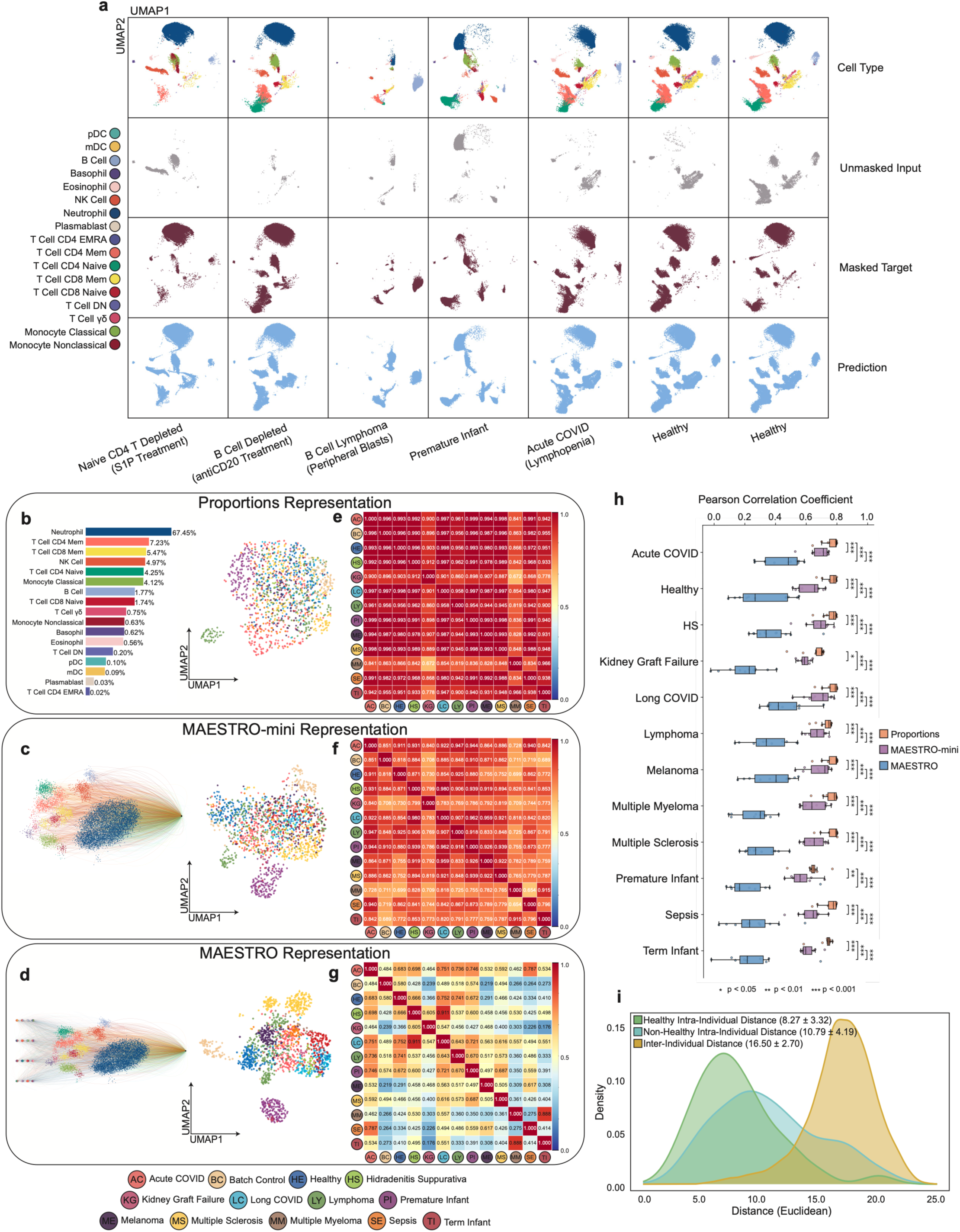
Self-supervised learning captures biologically meaningful immune structure. **a.** Population-level masked reconstruction across diverse immune states. For each sample: annotated cell populations (top), unmasked input cells, masked target cells, and MAESTRO predictions for masked cells, visualized in a shared two-dimensional embedding. Examples span treatment-induced depletion (S1P receptor modulator, anti-CD20), B cell lymphoma with peripheral blasts, premature infants, acute COVID-19 with lymphopenia, and healthy donors. **b.** Proportions Representation: two- dimensional projection of gated cell-type proportion vectors colored by diagnosis, with an example 17- dimensional composition (right). **c.** MAESTRO-mini representation encoding per-cell features without inter-cellular attention, shown as a two-dimensional projection colored by diagnosis. **d.** Full MAESTRO representation incorporating inter-cellular attention, shown as a two-dimensional projection colored by diagnosis. **e–g.** Pairwise Pearson correlation matrices between diagnoses computed from Proportions Representation (**e**) MAESTRO-mini (**f**) and MAESTRO embeddings (**g**) showing progressive reduction in cross-disease correlation with richer representations. **h.** Per-diagnosis distributions of pairwise correlation coefficients for Proportions Representation (orange), MAESTRO-mini (purple), and MAESTRO (blue), demonstrating reduced inter-disease similarity in the MAESTRO embedding space. Box plots show median (center line), interquartile range (box), and 2.5^th^–97.5^th^ percentiles (whiskers). Wilcoxon signed-rank test, adjusted *p < 0.05, **p < 0.01, ***p < 0.001. **i.** Kernel density estimates of Euclidean distances between MAESTRO embeddings for longitudinal samples from the same healthy individual (yellow), the same non-healthy individual (green), and between different individuals (blue), demonstrating stable person-specific immune fingerprints.

We next assessed whether MAESTRO embeddings captured immune system structure beyond conventional immune profiling approaches. We compared three representations of immune landscapes: conventional cell type proportions (Proportions Representation; **Fig. 2b**), an encoding where each cell is processed independently without modeling relationships between cells (MAESTRO- mini; **Fig. 2c**), and the full MAESTRO architecture with inter-cellular attention where the model learns relationships between cells in a sample (MAESTRO, **Fig. 2d**). UMAP projections revealed a gradient of resolution, with Proportions Representation showing minimal separation by disease, MAESTRO-mini showing modest improvement, and full MAESTRO clearly distinguishing different health and disease states. This organization was not driven by demographic covariates (**Supplementary Fig. 4**), and clustering of batch control samples indicated that batch effects did not explain the observed structure (**Fig. 2d**), nor could it be explained by cell type proportions alone (**Supplementary Fig. 5**).

Pairwise correlation analysis further quantified the differences between the three approaches (**Fig. 2e-g**). Proportions Representation yielded high cross-disease correlation exceeding 0.85 for most condition pairs, indicating poor capacity to distinguish disease-specific immune signatures (**Fig. 2e**). This low resolution reflected the dominance of shared major immune lineages when cell type proportions were averaged within each diagnosis because median proportions show only modest compositional shifts between diseases (**Supplementary Fig. 6a**). Increasing the number of cell types for this approach from 17 to 35 populations did not improve resolution (**Supplementary Fig. 6b**). MAESTRO-mini reduced cross-disease correlation relative to the Proportions Representation but retained substantial overlap between conditions (**Fig. 2f**). Full MAESTRO embeddings produced a correlation structure where disease-specific immune signatures were evident and cross-disease similarities were reduced (**Fig. 2g**). MAESTRO embeddings also preserved biologically meaningful relationships between related conditions. Premature and term infants showed high correlation (0.888) while diverging strongly from adult conditions. Acute COVID-19 and sepsis shared high correlation (0.787), consistent with the sepsis-like presentation observed in many hospitalized acute COVID-19 patients (**Fig. 2g**). HS and long COVID showed similarity to each other and to healthy controls, suggesting less pronounced immune perturbation. Aggregating the correlation analysis demonstrated that the Proportions Representations showed uniformly poor ability to distinguish diseases (high similarity across diseases), MAESTRO-mini showed intermediate performance, whereas full MAESTRO achieved significantly better ability to distinguish diseases (lower inter-disease correlation across all categories; Wilcoxon signed-rank test, adjusted p < 0.001 for all comparisons; **Fig. 2h**). These results demonstrated that MAESTRO embeddings resolved disease-specific immune architecture and outperformed other approaches for defining immune landscapes.

Finally, we tested whether MAESTRO was able to recover stable, subject- or patient-specific immune signatures without explicit training on individual identity. Prior work has shown that there is greater immune system variation between people than within the same individual sampled at different timepoints (22). We computed Euclidean distances between MAESTRO embeddings within the same person sampled at different times compared to between different individuals (**Fig. 2i**). Across cohorts, inter-person distances centered at 16.50 ± 2.70, whereas intra-person distances for healthy individuals centered at 8.27 ± 3.32, indicating that the same healthy individual sampled across time produced representations approximately half as distant as those from two different people. Individuals with disease showed higher intra-person distances (10.79 ± 4.19), consistent with pathological processes perturbing immune architecture over time, but these individuals remained more similar to themselves than to other individuals. These results confirm that MAESTRO embeddings capture stable individual-level immune fingerprints that persist through both health and disease, providing a quantitative basis for tracking immune states within individuals over time.

### MAESTRO enables sample-efficient disease classification with interpretable cellular attention

We next tested whether MAESTRO embeddings contained sufficient information to support disease prediction. The full MAESTRO embeddings organized immune states into more meaningful structure than conventional representations. We, therefore, subjected these embeddings to a more stringent test. We trained a multi-class logistic regression classifier on MAESTRO embeddings, repeating the experiment five times with a different held-out test set (five-fold cross-validation) across 13 disease categories (**Fig. 3a**). In other words, a blood sample of unknown clinical origin could be classified into any of these 13 disease categories based solely on its immune embedding. The confusion matrix in **Fig. 3a** reveals a strong diagonal structure where most samples were classified correctly at rates exceeding 70%. Premature infants (98.1%), melanoma (97.4%), and multiple myeloma (95.9%) achieved near-perfect classification. ANCA vasculitis (83.8%), acute COVID-19 (78.3%), and lymphoma (87.0%) showed high accuracy with minimal confusion across unrelated conditions. Healthy control samples classified at 80.3% accuracy, with modest misclassification to diseases associated with less perturbed immune states.

**Figure 3:**
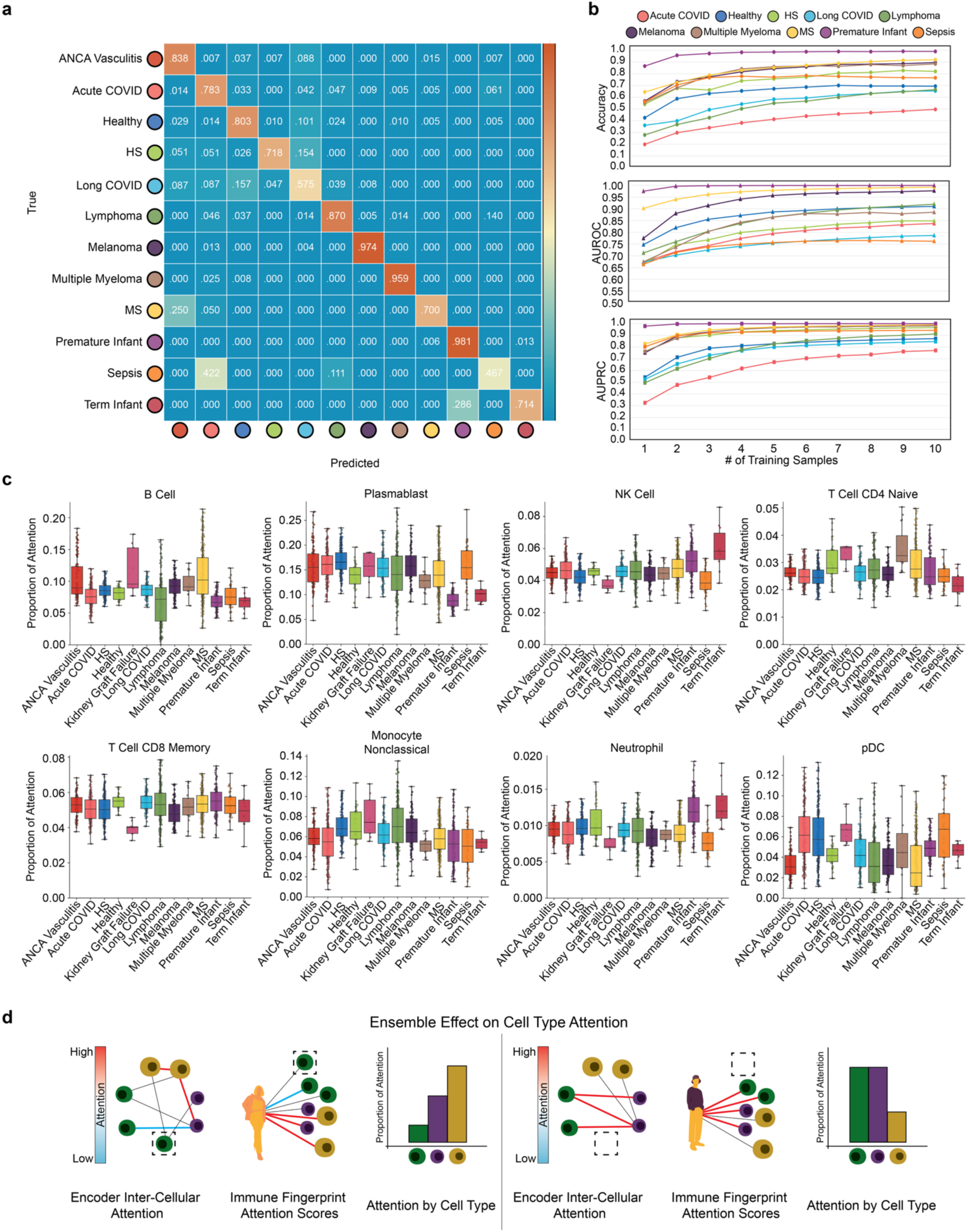
MAESTRO enables sample-efficient disease classification with interpretable cellular attention. **a.** Multi-class classification from frozen MAESTRO embeddings using logistic regression with five-fold cross-validation. Confusion matrix shows prediction performance across 13 diagnostic categories (rows, true labels; columns, predicted labels); values indicate the fraction of samples assigned to each predicted class. **b.** Few-shot classification performance using progressively larger labeled training sets. Accuracy (top), AUROC (middle), and AUPRC (bottom) as a function of training samples per class, with error bars indicating standard deviation across folds. **c.** Attention allocation across representative cell types. Box plots show the proportion of total pooling attention assigned to each gated population across diagnostic categories, computed by aggregating per-cell attention weights within each population. Boxes show median (center line), interquartile range (box), and 2.5^th^–97.5^th^ percentiles (whiskers); points denote individual samples. **d.** Schematic of hierarchical feature encoding in MAESTRO. Early encoder layers represent marker expression patterns corresponding to canonical cell types, while deeper layers encode inter-cellular attention across all cells in a sample. The final attention score assigned to each cell reflects both its own marker expression and the broader immunological context in which it appears, meaning that cells sharing the same canonical label can receive different attention weights depending on the composition and state of the surrounding immune landscape.

Certain misclassification patterns reflected biological relationships. MS samples misclassified most frequently as ANCA vasculitis (25.0%), both autoimmune conditions with shared inflammatory features (44). Sepsis showed overlap with acute COVID-19 (42.2%), consistent with related acute inflammatory signatures (45). Term infants misclassified partially as premature infants (28.6%), reflecting the developmental continuum of neonatal immunity. Long COVID had the most distributed misclassification, consistent with the heterogeneous clinical presentation and a still unclear but likely complex immunological basis (46). These biologically coherent patterns suggested that MAESTRO embeddings captured immunological similarity that cuts across diagnostic boundaries. Moreover, because this classifier test was intentionally simple, strong performance cannot be attributed to the classifier itself. This accuracy in disease identification must originate from the information contained in the embeddings of immune system structure.

An advantage of pretrained representations is sample efficiency. Because the encoder has already learned generalizable immune structure from large-scale unannotated data, downstream classifiers require fewer labeled samples to achieve strong performance (47–49). We, therefore, evaluated “few-shot” classification performance by training logistic regression classifiers on progressively larger subsets of samples of known identity for the disease prediction task (**Fig. 3b**). With only a single training example per class, most diseases achieved above-chance accuracy, and performance for downstream prediction improved rapidly with additional samples. By ten training examples, accuracy, AUROC (measuring overall discrimination between classes), and AUPRC (measuring precision across sensitivity thresholds) approached asymptotic values for most diseases (**Fig. 3b**). Melanoma and premature infants achieved maximum performance with minimal data, whereas more heterogeneous diseases like sepsis and long COVID benefited from additional training samples. This efficiency of disease prediction from small numbers of samples was achieved because of the size of the corpus of training data (∼1800 samples; ∼ million cells) and robustness of the MAESTRO model. Indeed, these observations suggest approaches using MAESTRO or similar models that could enable clinical applications where cohorts may be small, allowing disease-specific classifiers to be trained on modest pilot datasets rather than requiring large-scale annotation efforts.

We next sought to understand which immune features drive the ability of MAESTRO representations to distinguish disease states. The pooling module (attention block in MAESTRO architecture) assigns attention weights to each cell, quantifying its contribution to the final sample embedding. To enable biological interpretation, we aggregated attention scores by canonically defined cell type (**Fig. 3c, Supplementary Fig. 7**).

Across diseases, several general principles emerged from these MAESTRO attention patterns. Neutrophils numerically dominated many samples (**Fig. 1m; Supplemental Fig. 7**) yet received consistently low attention weights (**Fig. 3c**), demonstrating that attention was not strictly a feature of cellular abundance. The model learned that rarer populations such as pDCs in acute COVID-19 or residual B cells in anti-CD20 treated patients carried more discriminative information for distinguishing immune states, highlighting the value of acquiring large numbers of cells per sample to ensure acquisition of rare cell type data (**Fig. 3b, 3c**).

Infant cohorts showed increased attention on neutrophils, NK cells, eosinophils, and gamma-delta T cells, with reduced attention on plasmablasts (**Fig. 3c**), consistent with the innate-biased, antigen-inexperienced immune architecture of early life (50–52). Gamma-delta T cells, which provide early immune protection before conventional T cell repertoires mature, received particularly high attention in premature infants (53). Kidney graft failure produced a distinct signature with elevated attention on naive CD4 T cells and reduced attention on CD8 memory T cells, double-negative T cells, and gamma-delta T cells (**Fig. 3c**), potentially reflecting immunosuppressive therapy effects on T cell populations. ANCA vasculitis showed elevated B cell attention, multiple myeloma showed elevated attention on z memory and naive CD4 T cells rather than B lineage cells, and sepsis directed attention toward CD4 memory T cells and plasmacytoid dendritic cells (**Fig. 3c**). Thus, MAESTRO captured and distinguished immune perturbations associated with disease including identifying many of known relevance and identifying others that may warrant future investigation.

One observation from these attention scores was that key immune populations appeared to serve as condition-agnostic anchors. Classical monocytes, nonclassical monocytes, and naive CD8 T cells maintained stable attention regardless of disease state (**Fig. 3c**), suggesting these populations provided baseline immune context rather than condition-discriminating features. The contrast between disease-variable and disease-stable attention further supports the conclusion that MAESTRO captures organized immune architecture rather than simply reflecting cell type abundance.

Although cell-type aggregation revealed where the model focused attention, this analysis understated the information captured. Deep neural networks learn hierarchical representations, where early layers encode superficial features while deeper layers capture increasingly abstract structure (54). For MAESTRO, early encoder layers likely represented marker expression patterns corresponding to canonical cell types, while deeper layers encoded inter-cellular attention across all cells in a sample, meaning that the final attention score assigned to any individual cell reflected not only a cell marker expression pattern but also the broader immunological context in which it appears (**Fig. 3d**). The attention scores aggregated by cell type in **Fig. 3c** therefore already encoded information about immune system composition and inter-population structure, not merely cell-type identity. Because inter-cellular attention operates at the level of individual cells rather than cell type categories, cells sharing the same label can receive substantially different attention weights. To quantify this within-type variation, we computed the coefficient of variation (CV) of attention scores among cells within each cell type per sample. Median CV across all samples and cell types was 0.617, with 67% of sample and cell type pairs exceeding 0.5 (**Supplementary Fig. 8a**), confirming that MAESTRO assigned substantially different weights to cells sharing the same label. This within-type variability was consistent across cell types (**Supplementary Fig. 8b**) and varied across disease conditions (**Supplementary Fig. 9**), suggesting that fine-grained discrimination among phenotypically similar cells may itself encode disease-relevant information beyond what cell-type aggregation captures.

### Learned representations encode temporal immune information

The results above demonstrate that MAESTRO learns complex immune features that can accurately distinguish static health and disease states. We next tested whether MAESTRO could also resolve temporal aspects of immune biology. We hypothesized that immune fingerprints encode information about both immune history and future potential. To test this idea, we performed two independent analyses: first, we examined whether MAESTRO representations retained signatures of past exposures, and second, we tested whether baseline immune profiles could predict future immune responses.

To address the first question, we evaluated whether MAESTRO could identify a previous immunological event. We analyzed a dataset of peripheral blood samples from infants who were exposed to HIV in utero but remained uninfected (HEU) versus unexposed (UE) control infants (**Fig. 4a**). Previous studies demonstrated that HIV exposure signatures persist at 6 months in the absence of overt infection, long after maternal antibodies have waned (55, 56) indicating that early life exposures reshape immune architecture in ways detectable months later. The question here is, if one did not know the HIV status of the mother, could the immunological signatures distinguish infants exposed versus unexposed to HIV in utero. We applied MAESTRO to the Flow Cytometry Critical Assessment of Population Identification Methods (FlowCAP)-IV challenge (57), a community-driven benchmark that enables direct comparison across computational cytometry methods on standardized biological questions. In this case the question was whether different methods could distinguish exposed versus unexposed infants based on blood cytometry profiles alone. This FlowCAP-IV dataset was collected with standard fluorescent cytometry, requiring retraining of MAESTRO (**Methods**) to accommodate fluorescence-based measurements and a distinct antibody panel compared to the CyTOF-based data above. Each infant blood sample was profiled using multiple immune stimulation conditions (unstimulated, PAM, LPS, PG, PIC, CPG, R848) to test whether previous HIV exposure signatures persist even in the presence of strong acute immune perturbations.

**Figure 4:**
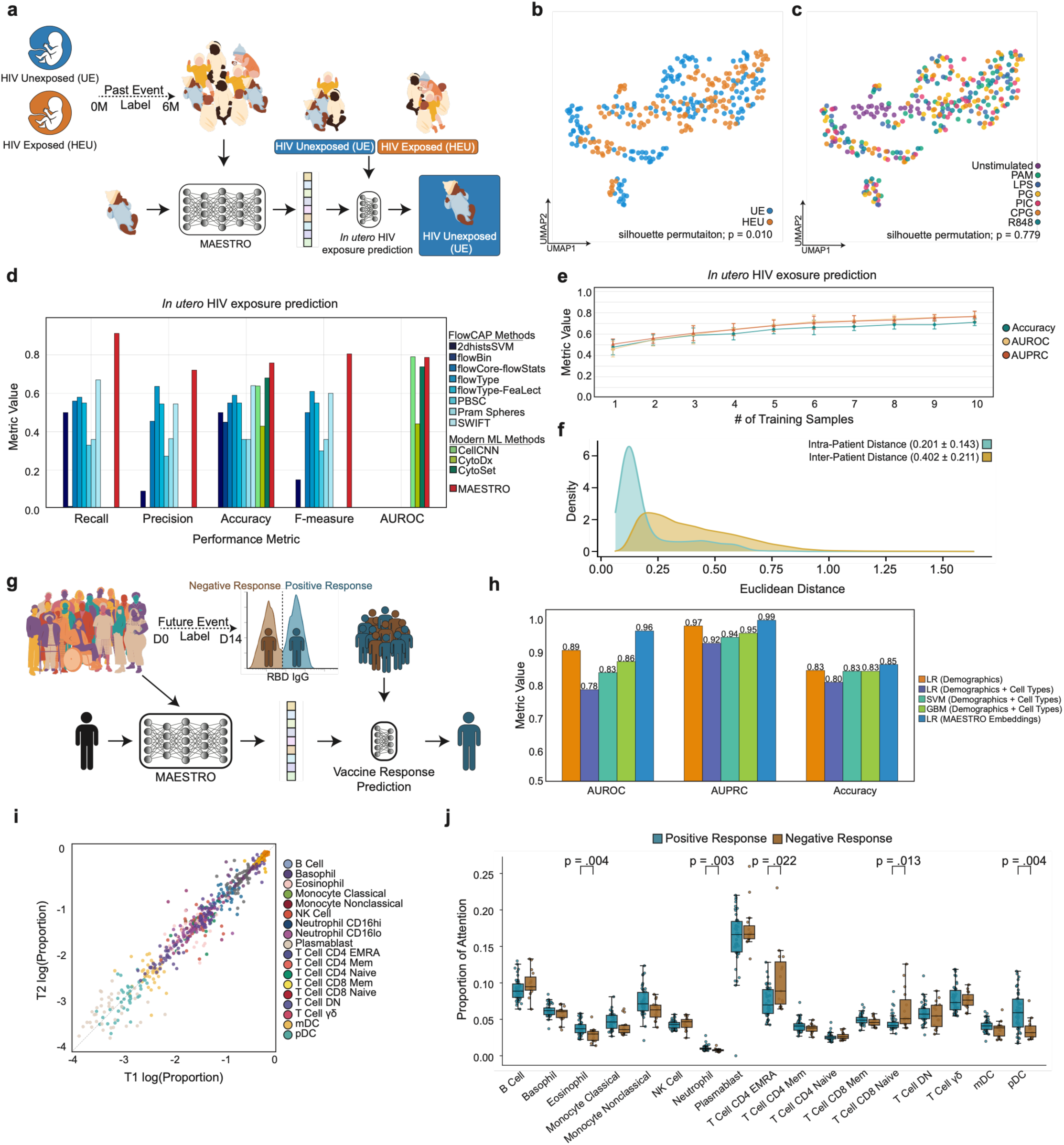
Learned representations encode temporal immune information spanning past exposures and future responses. **a.** Schematic of the in utero HIV exposure classification task: 6-month postnatal blood samples from HIV-exposed uninfected (HEU) and unexposed (UE) infants are encoded by MAESTRO and classified by exposure status. **b.** UMAP of MAESTRO embeddings for the FlowCAP-IV cohort colored by exposure status (HEU versus UE). **c,** Same embedding colored by stimulation condition (unstimulated, PAM, LPS, PG, PIC, CPG, R848). **d.** Benchmarking of HEU detection: recall, precision, accuracy, F- measure, and AUROC for MAESTRO compared with FlowCAP-IV challenge methods and recent cytometry deep learning baselines. **e.** Few-shot HEU classification performance (accuracy, AUROC, AUPRC) as a function of labeled training samples per class. Error bars indicate variability across resampling runs. **f.** Kernel density estimates of Euclidean distances between MAESTRO embeddings from the same infant across stimulation conditions (intra-patient) versus between different infants (inter- patient). **g.** Schematic of the vaccine response prediction task: pre-vaccination (D0) immune profiles are used to predict post-vaccination antibody response (RBD IgG). **h.** Vaccine response prediction performance (AUROC, AUPRC, accuracy) for logistic regression on demographics alone, demographics plus cell-type proportions across multiple classifiers, and logistic regression on MAESTRO embeddings. **i.** Comparison of log-transformed cell-type proportions before (T1) and after (T2) vaccination, with each point representing a cell population. **j.** Pooling attention by cell type for baseline samples stratified by future vaccine response. Box plots show median (center line), interquartile range (box), and 2.5^th^–97.5^th^ percentiles (whiskers); points denote individual samples. P- values from two-sided Mann–Whitney U tests.

UMAP projection of MAESTRO embeddings showed modest separation between HEU and UE infants (silhouette permutation test, p = 0.010) (**Fig. 4b**), though visual separation in compressed projections is an imperfect proxy for discrimination. When colored by stimulation condition, samples intermingled throughout the embedding space (silhouette permutation test, p = 0.779) (**Fig. 4c**), confirming that acute stimuli did not drive clustering. Instead, samples from the same infant clustered together across stimulation conditions (**Supplementary Fig. 12**), demonstrating that individual immune identity dominated over transient activation in MAESTRO representations.

To directly test whether previous HIV exposure was distinguishable in MAESTRO representations, we benchmarked against other approaches in the FlowCAP-IV challenge. MAESTRO correctly identified 91.2% of truly HIV-exposed infants, missing very few cases (low false negatives; recall), and 72.1% of infants flagged as exposed were genuinely HI-exposed, meaning most flags were not false alarms (low false positives; precision). Across all infants regardless of exposure status, 75.8% were correctly classified (accuracy) (**Fig. 4d**). Combining these measures into a single score that balances the ability to detect exposed infants against the rate of false positives produced an F-measure of 0.805 and an AUROC of 0.787, which quantifies how well the model separated exposed from unexposed infants overall. Using the same training and testing samples, MAESTRO outperformed the eight original methods developed specifically for this challenge. These previous methods were largely based on manual feature engineering and conventional statistical classifiers that struggled to identify robust correlates of exposure. MAESTRO also achieved superior or comparable performance against more recent deep learning approaches including CellCNN (58), CytoDx (59), and CytoSet (60), which are methods that were specifically designed for this task (supervised). Few-shot experiments further demonstrated data efficiency for HEU detection (**Fig. 4e**). Accuracy reached approximately 0.65 with five samples and plateaued around 0.71 by ten samples, while AUROC climbed from 0.68 to over 0.77, indicating that pretrained embeddings organize HEU and UE samples into separable structure requiring only a few labeled examples to define decision boundaries. These results indicate that general-purpose self-supervised pretraining efficiently captured subtle immune remodeling from past exposures outperforming previous specialized approaches designed specifically for this task.

To quantify the stability of individual immune landscapes across stimulation conditions, we computed Euclidean distances between MAESTRO embeddings within the same infant across all stimulations versus between different infants (**Fig. 4f**). Intra-patient distances centered at 0.201 ± 0.143, whereas inter-patient distances centered at 0.402 ± 0.211. This consistent two-fold difference between intra- and inter-subject immune landscape maps across independent datasets, suggesting a fundamental property of immune organization: individual identity accounts for roughly half the information captured by MAESTRO embeddings. This feature of MAESTRO enables exposure classification because individual immune fingerprints, while dominant, do not obscure the subtler exposure-driven structure in the embedding space.

To further explore whether MAESTRO-derived baseline immune profiles could predict future responses, we used samples obtained from individuals before COVID-19 vaccination to test whether MAESTRO could predict vaccine responses. We trained classifiers to predict antibody response measured by IgG RBD ELISA at day 14 post-vaccination, where a positive response was defined as IgG concentration above 0.48ug/mL and negative response below 0.48ug/mL (**Fig. 4g**).

Using only pre-vaccination immune profiles when training the model, MAESTRO embeddings achieved strong discrimination of positive versus negative vaccination outcomes. The model correctly classified 85% of individuals (accuracy) and separated responders from non-responders with an AUROC of 0.96, indicating near-perfect discrimination (**Fig. 4h**). Among predicted responders, AUPRC of 0.99 confirmed that the model identified true responders with high reliability. MAESTRO outperformed conventional approaches including logistic regression on demographics alone (AUROC 0.89), logistic regression and support vector machine (SVM) on demographics combined with cell type proportions (AUROC 0.78 and 0.83 respectively), and gradient boosted machines (GBM) on the same features (AUROC 0.86) (**Fig. 4h**). UMAP projection of MAESTRO embeddings showed partial separation between positive and negative responders at baseline, before vaccination (**Supplementary Fig. 13**).

Cell type proportions between pre-vaccination (T1) and post-vaccination (T2) timepoints correlated strongly across all populations, with points clustering tightly along the diagonal (**Fig. 4i**). Vaccination did not substantially alter overall immune composition, consistent with the notion that vaccine responses are mediated by relatively rare cells in the blood while most circulating immune cells are uninvolved bystanders. This overall immune stability suggested that the predictive performance of MAESTRO derives not from changes in frequency of major immune populations but from subtler features of immune organization that distinguish future responders from non-responders at baseline.

Analysis of attention scores revealed cellular populations that may contribute to vaccine response prediction (**Fig. 4j**). Eosinophils, neutrophils, EMRA CD4 T cells, naive CD8 T cells, and pDCs showed significantly different attention weights between responders and non-responders (**Fig. 4j**). Eosinophils, neutrophils, and pDCs received elevated attention in individuals who mounted positive responses, whereas T cells received higher attention in future non-responders. The elevated attention on pDCs in responders is consistent with the role of these cells as early sentinels for mRNA vaccines, where they act as the primary source of Type I interferons required to initiate a robust humoral response (55, 56). Similarly, although neutrophils and eosinophils are traditionally viewed as end-stage effectors, recent work identifies these cells as key regulators that shape early antibody production through rapid cytokine release and plasmacell support (61). Conversely, the high attention on T cell subsets in non-responders, specifically EMRA populations, may capture a signature of immunosenescence (62). These attention differences, derived from baseline samples before antigen exposure, suggest that the capacity for productive vaccine responses is encoded in pre-existing immune architecture. Although not all attention patterns will point to causal features, some insights may provoke future direct biological interrogation.

### Precision immunotherapy using immune profiles

We next tested whether this MAESTRO framework could be translated to the clinical challenge of predicting immunotherapy responses in cancer patients. We tested this idea in patients with pancreatic ductal adenocarcinoma (PDAC), a malignancy with few therapeutic options and a five-year survival rate for advanced disease of < 5% (63, 64). We applied MAESTRO to samples from the PRINCE trial, a randomized phase 2 trial evaluating the efficacy of nivolumab (anti-PD-1) and/or sotigalimab (anti-CD40 agonistic antibody) with gemcitabine/nab-paclitaxel chemotherapy in previously untreated metastatic PDAC patients (**Fig. 5a**) (65). The primary endpoint of 1-year overall survival (OS) was met for nivolumab/chemotherapy (57.7%) but not for sotigalimab/chemotherapy (48.1%) or the triple combination (41.3%) (65). Previous traditional flow cytometric analysis of longitudinal patient samples from this trial identified distinct immune signatures associated with survival for each treatment arm. However, it has not been possible to prospectively stratify patients onto the optimal treatment arm prior to therapy, a major unmet need for cancer and other diseases. Thus, using this PRINCE trial data, we asked whether MAESTRO could predict treatment response from baseline immune profiles, enabling personalized therapy selection before treatment begins.

**Figure 5:**
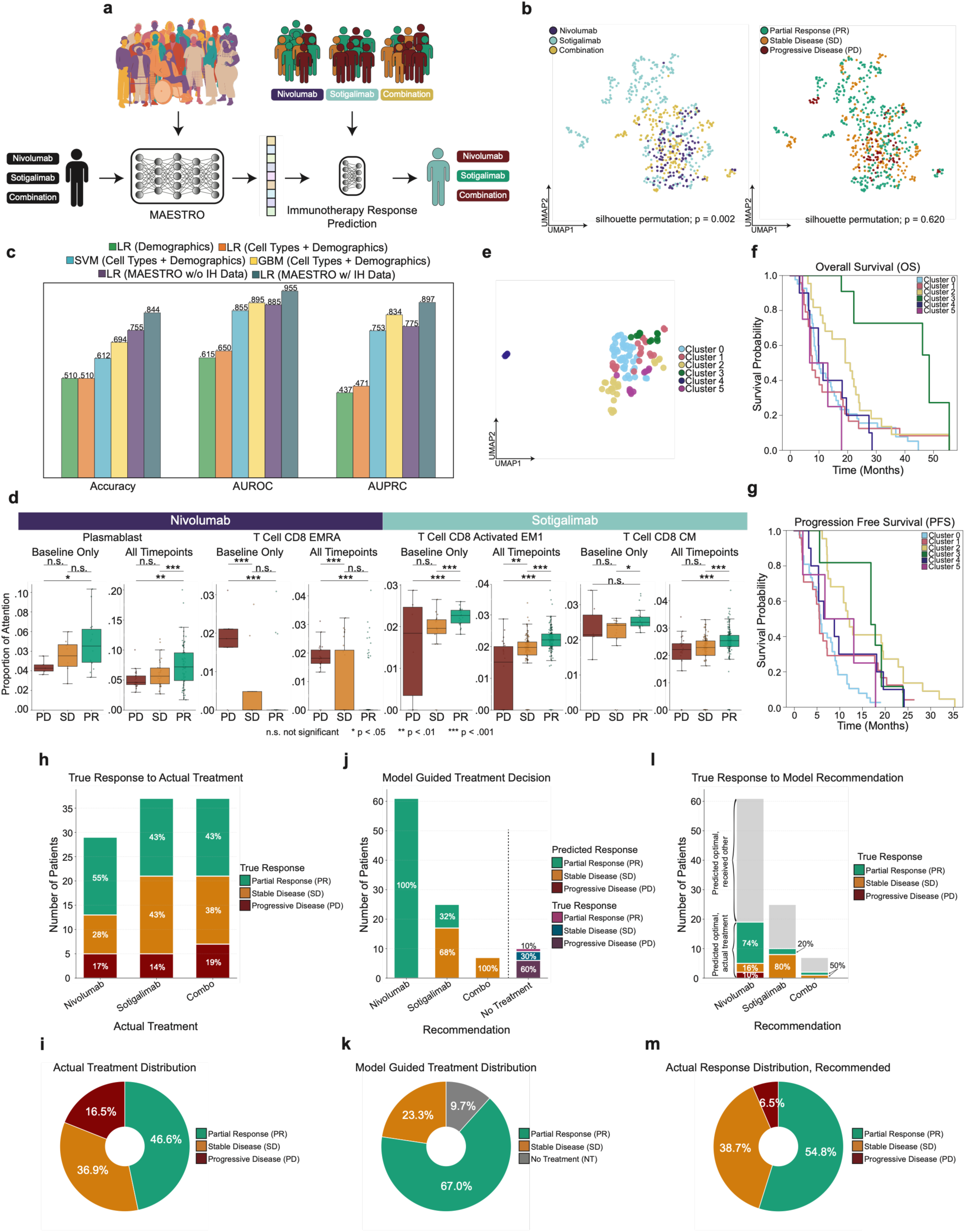
Precision immunotherapy using baseline immune fingerprints in metastatic PDAC. **a.** Study and modeling framework for the PRINCE trial. Pre-treatment peripheral blood samples are encoded by MAESTRO, and embeddings are input to arm-specific models predicting response to nivolumab, sotigalimab, or combination therapy. **b.** UMAP of MAESTRO embeddings for PRINCE samples colored by treatment arm (left) and clinical response (right; PR, partial response; SD, stable disease; PD, progressive disease). **c.** Leave-one-out cross-validation performance for treatment response prediction. Accuracy, AUROC, and AUPRC are shown for demographics alone, demographics plus cell-type proportions across multiple classifiers, and logistic regression on MAESTRO embeddings pretrained with PRINCE data only or with the full pretraining Immune Health (IH) corpus (**Fig. 1**). **d.** Treatment-specific cell type attention signatures associated with clinical response categories. Box plots show pooling attention for selected cell populations stratified by response category for the nivolumab or sotigalimab arms, evaluated at baseline only or across all timepoints. Boxes show median (center line), interquartile range (box), and 2.5^th^–97.5^th^ percentiles (whiskers); points denote individual samples. P-values from two-sided Mann–Whitney U tests; *p < 0.05, **p < 0.01, ***p < 0.001. **e.** UMAP of baseline (C1D1) MAESTRO embeddings colored by cluster assignment. **f,g.** Kaplan–Meier survival curves for patient clusters identified by hierarchical clustering of baseline MAESTRO embeddings: overall survival (**f**) and progression-free survival (**g**). **h.** Distribution of true clinical responses (PR, SD, PD) within each treatment arm. **i.** Aggregate response distribution across all patients using actual trial randomization. **j.** Model-guided treatment assignment: each patient is assigned to the arm with highest predicted response probability based on baseline embeddings; patients predicted to be unlikely to benefit from any arm are assigned to a no-treatment category. Stacked bars show predicted response composition per arm and true outcomes for the no-treatment group. **k.** Aggregate predicted outcome distribution using model-guided assignment. **l.** True clinical responses for patients who received the MAESTRO-recommended treatment; inset shows the fraction of patients who received versus did not receive their predicted optimal arm. **m.** Aggregate response distribution among patients who received their MAESTRO-recommended treatment, showing 54.8% PR, 38.7% SD, and 6.5% PD.

MAESTRO was pretrained on diverse cytometry datasets spanning autoimmunity, infection, malignancy, and healthy controls from the cohorts in **Fig. 1**, combined with pre-treatment (C1D1) samples from PRINCE (**Fig. 5a**). For a new PDAC patient, MAESTRO generated an embedding from their pre-treatment cytometry profile, which was then fed into a task-specific model trained on PRINCE data to predict response to each therapy. Predictions were made from baseline immune architecture alone, before patients received any intervention. This design tested two hypotheses: whether the immune fingerprint of each patient prior to treatment contained sufficient information to predict therapeutic outcomes, and whether immune structure learned from unrelated conditions transferred to cancer immunotherapy prediction. Such knowledge transfer would enable predictive insights even for clinical trials where limited sample sizes would not be sufficient for training on their own.

MAESTRO embeddings organized PDAC samples into structured representations when visualized by treatment arm (**Fig. 5b, left**) (Silhouette permutation test, p = 0.003), though response categories did not show significant spatial clustering (**Fig. 5b, right**) (Silhouette permutation test, p = 0.620). Although MAESTRO was pretrained only on baseline PRINCE timepoints, these projections included all timepoints from the trial, suggesting that the learned immune structure generalizes across treatment-induced perturbations.

We next evaluated predictive performance of treatment response using a leave one out cross validation strategy. The training set included samples with all available timepoints while the left-out test sample was evaluated using only the baseline timepoint. Logistic regression on MAESTRO embeddings outperformed conventional machine learning approaches for predicting treatment response (**Fig. 5c**). Models using demographics alone (age and sex) achieved 0.510 accuracy, 0.615 AUROC, and 0.437 AUPRC (**Fig. 5c**). The addition of cell type proportions provided marginal improvement regardless of model complexity where logistic regression reached 0.510 accuracy, 0.650 AUROC, and 0.471 AUPRC while SVM achieved 0.612 accuracy, 0.855 AUROC, and 0.753 AUPROC (**Fig. 5c**). GBM performed slightly better with 0.694 accuracy, 0.895 AUROC, and 0.834 AUPRC (**Fig. 5c**). MAESTRO embeddings generated by pretraining MAESTRO with only PRINCE trial data (**Fig. 1**) achieved 0.755 accuracy and 0.885 AUROC with 0.775 AUPRC using simple logistic regression (**Fig. 5c**). Including the diverse cohorts from **Fig. 1** in pretraining raised performance to 0.844 accuracy with 0.955 AUROC and 0.897 AUPRC (**Fig. 5c**). This improvement from incorporating unrelated diseases highlights that learning general immune system structure improves cancer-specific prediction. The performance gap between MAESTRO and conventional approaches confirms that baseline immune fingerprints contain rich predictive information not captured by summary cell proportions.

Interrogating the MAESTRO attention patterns revealed treatment-specific cellular signatures distinguishing responders from non-responders (**Fig. 5d**). With nivolumab treatment, Plasmablasts received significantly elevated attention in partial responders compared to progressive disease patients (p = 0.019 baseline, p = 0.005 all timepoints) (**Fig. 5d**). CD8 EMRA T cells showed higher attention in progressive disease patients compared to partial response and stable disease patients at baseline (p < 0.001) (**Fig. 5d**). This pattern remained when using all timepoints for CD8 EMRA T cell attention across response groups. With sotigalimab treatment, CD8 activated EM1 T cells received elevated attention in partial response patients compared to progressive disease (p < 0.001 baseline, p < 0.001 all timepoints), and CD8 central memory T cells showed higher attention in partial responder patients at baseline (p = 0.022) with this pattern strengthening when all timepoints were included (p < 0.001 for partial response versus either stable disease or progressive disease) (**Fig. 5d**). These patterns were largely consistent between baseline-only and all-timepoint analyses (**Fig. 5d**), suggesting that the immune features distinguishing response groups were present before treatment and persisted through therapy. It is likely that deeper layers of attention focused on more complex or abstract immune features exist (54) that will warrant future investigation.

Unsupervised hierarchical clustering of baseline MAESTRO embeddings identified six patient subgroups with distinct survival trajectories (**Fig. 5e-g**). These clusters occupied distinct regions of the MAESTRO embedding space (**Fig. 5e**), confirming that the survival-associated subgroups correspond to distinguishable immune architectures. For OS, Cluster 3 patients experienced superior outcomes, with an approximate survival probability of 0.7 at ∼45 months whereas for most other patient clusters this survival probability declined below 0.2 by 30 months (**Fig. 5f**). Patients in clusters 0, 1, 4, and 5 had poor survival early, whereas Cluster 2 patients had an intermediate trajectory. For progression-free survival (PFS), the patterns differed slightly: Patients in Cluster 2 and Cluster 3 experienced the longest disease control, with progression free survival probability above 0.3 at 18 months (**Fig. 5g**). In contrast, patients in Clusters 0, 1, 4, and 5 had poor PFS in the first 10 months (**Fig. 5g**). Thus, MAESTRO was able to discover immune architectures from baseline samples that then separated patients by clinical trajectory.

The preceding analyses demonstrated that baseline immune fingerprints could predict treatment response in an immunotherapy trial. However, a major clinical goal is matching each patient to their optimal therapy based on individual immune fingerprints prior to therapy. We next asked whether MAESTRO-guided treatment assignment could improve treatment outcomes compared to the randomization approach used in the PRINCE trial (**Fig. 5h-m**). Using randomization, the trial achieved overall 46.6% PR, 36.9% SD, and 16.5% PD, with response rates varying across treatment arms (**Fig. 5h-i**).

MAESTRO assigned each patient to the therapy arm with highest predicted response probability based on their baseline immune fingerprint (**Fig. 5j**). Using this framework, 61 patients were assigned to nivolumab, 25 to sotigalimab, and 7 to combination therapy. MAESTRO also identified 10 patients for whom no treatment was predicted to be effective. Among these patients, 60% indeed experienced progressive disease regardless of which treatment arm they received, 30% achieved a SD response, and 10% a partial response. This capacity to identify patients unlikely to benefit from available therapies may itself be clinically useful, potentially sparing patients from ineffective treatment and enabling assignment to alternative trials. Overall, the resulting predicted trial outcome distribution shifted substantially to 67.0% PR and 23.3% SD, compared to 46.6% PR using randomization (**Fig. 5k**).

Given that these are predictions, we next asked what happened for the patients who actually received the treatment MAESTRO predicted as optimal. Among patients who received MAESTRO- recommended nivolumab, 74% achieved PR, 16% SD, and 10% PD (**Fig. 5l**). Sotigalimab resulted in 20% PR and 80% SD with no progressive disease, while combination therapy yielded 50% PR and 50% SD, though numbers for combination therapy were low (**Fig. 5l**). Combined, 54.8% achieved partial response, 38.7% stable disease, and 6.5% progressive disease, with an overall 93.5% benefitting from treatment compared to 83.5% overall benefit and 46.6% PR using randomization (**Fig. 5m**). These findings suggest that baseline immune architecture may guide therapy selection in a way that improves response rates. Although prospective validation is required, these results have implications beyond clinical trials to broader clinical settings where immune-modifying treatments are used.

## Discussion

Here, we developed MAESTRO, a self-supervised deep learning framework that transforms high-dimensional cytometry data into compact representations of individual immune health. Pretrained on over 418 million immune cells across 1,792 samples spanning 13 clinical diagnoses, MAESTRO learns immune fingerprints that resolve disease-specific immune architecture beyond what conventional cell type proportions capture. These fingerprints are stable within individuals yet sensitive to disease-driven perturbations, providing a quantitative basis for comparing immune states across individuals and time. MAESTRO embeddings support sample-efficient disease classification through simple models, encode temporal information spanning past immune exposures and future response potential, and enable prospective prediction of immunotherapy responses in metastatic PDAC. Together, these results establish a foundation for translating immune complexity into clinically actionable representations for precision immunology.

A central insight from this work is that the immune system stores information about past experiences and future potential in the overall landscape architecture connecting individual cells, in addition to the cell intrinsic changes associated with innate or adaptive immune memory. Such immune network information storage parallels how neural circuits store memories through patterns of connectivity rather than only the biology of individual neurons (66). MAESTRO was designed to capture this relational immune structure by encoding information from hundreds of thousands of cells per individual. We prioritized the breadth of cellular sampling to capture inter-population relationships and ensemble network properties, including from underrepresented cell populations in blood, over approaches that might provide more depth of information for individual cells (e.g. single cell genomics). The latter approaches are well-suited for characterizing individual cells and cell-autonomous programs (i.e. models of a cell), but may scale less well for defining network architectures for large numbers of cells and may under sample less abundant cell types where much of the network variance exists in the human immune system. High dimensional immune profiling has been a powerful approach to annotate immune changes (37). Although these approaches have revealed much about human immunology, analyses are often centered on changes in cell type proportions or activation states. This type of analysis can provide insights about a known cell type or identify major perturbations to the frequencies of key responding cells, but often has difficulty distinguishing more complex disease states and trajectories. Moreover, individual disease cohorts are often too small for more integrated analytical approaches. To overcome some of these challenges MAESTRO embeddings learned from a large number of samples across diverse disease states and revealed information contained in this immune network ensemble. These embeddings were able to reveal underlying features residing in inter-population dependencies and deeper features of the immune network, capturing not only information about the unique immune fingerprint of individuals, but also about their immune and disease trajectories. The consistent two-fold ratio between intra- and inter-individual embedding distances across independent datasets and experimental conditions further supports this concept. Thus, applying MAESTRO revealed individual immune identity encoded in relational structure between cell populations producing a stable immune architectural signature shaped by genetics and cumulative immune exposure history that persists through health and disease.

The temporal dimensions encoded in MAESTRO embeddings have implications for how immune information is stored and accessed. MAESTRO detected signatures of in utero HIV exposure months after the exposure itself, indicating that past immunological events leave durable imprints on immune network architecture that persist beyond the resolution of the initiating perturbation. Pre-vaccination immune profiles predicted future antibody responses with high discrimination, suggesting that the capacity for future productive immune responses is encoded in baseline immune organization before antigen encounter. These observations are consistent with the concept that the immune system functions as a biological record of cumulative exposures while simultaneously encoding future response potential. The PRINCE trial analyses extend this principle to clinical decision-making, where baseline immune fingerprints predicted treatment response to immunotherapy and MAESTRO-guided treatment assignment improved predicted response rates from 46.6% to 67.0% for partial response. Among patients who received their MAESTRO-recommended therapy, 93.5% experienced clinical benefit compared to 83.5% using the original trial randomization approach. These results support the feasibility of immune-guided treatment selection, where pre-treatment immune architecture informs therapeutic matching. Thus, further scaling approaches like MAESTRO to including more data and more diverse immune states and treatment response patterns may be a powerful approach to develop precision immunotherapy across diseases.

Several limitations of this work warrant consideration. The training corpus, while large relative to existing single-cell foundation models, is derived primarily from peripheral blood profiled by mass cytometry with a 30-marker panel. Extension to tissue immune compartments, alternative or larger cytometry panels, and integration with other data modalities such as antibody serology, proteomics, and transcriptomics will likely capture additional features of immune organization. The attention-based interpretability analyses provide biological insight focused on discreet cell types that may be of interest. However, the deeper learned features that likely drive much of the discriminative capacity of MAESTRO remain difficult to map onto conventional immunological categories and future work will be necessary to understand how information is stored in the immune ensembles of these networks. The PRINCE trial counterfactual analysis, while supported by patients who received their MAESTRO-recommended therapy, requires prospective validation in a trial designed for immune-guided randomization. Similarly, the two-stage framework assumes that general immune structure learned from diverse conditions transfers to new clinical contexts, an assumption that held for PDAC but requires testing across additional diseases, treatment modalities, and patient populations. Finally, although longitudinal samples exist in the current corpus of data, the current framework operates on single timepoint snapshots. Incorporating longitudinal sampling directly into the model architecture could further improve the capacity to track immune dynamics and predict clinical trajectories.

The development of MAESTRO establishes a reusable foundation for precision immunology by converting high-dimensional immune complexity into structured, clinically actionable representations. This framework addresses a gap between the recognized complexity of the human immune system and the quantitative tools available to leverage that complexity for patient care. This approach capitalizes on the concept that information is stored not only in changes to individual immune cells over time, but also in changes to the overall immune network, in the ensemble structures of large numbers of immune cells. As cytometry datasets grow in scale and diversity and as immune-modulating therapies expand across diverse diseases, the ability to encode individual immune architecture into compact representations that predict disease trajectories and therapeutic responses has wide-ranging translational potential. More broadly, these results support a vision of precision medicine grounded not solely in static genetic information but in the dynamic, experience-shaped architecture of the immune system.

## Methods

### 1. Study cohorts and sample collection

#### 1.1 Pretraining cohort assembly

Peripheral blood samples were collected at the University of Pennsylvania and the surrounding area. All participants or their surrogates provided informed consent in accordance with protocols approved by the regional ethical research boards and the Declaration of Helsinki. Information about individual cohorts can be found below. Detailed information can be found in **Supplementary Table 1**. Longitudinal sampling information can be found in **Supplementary Table 2**.

##### ANCA Vasculitis

Subjects with antineutrophil cytoplasmic antibody (ANCA)-associated vasculitis (AAV) were consented and enrolled at the University of Pennsylvania Vasculitis Center during routine care. Individuals with the following AAV subtypes were analyzed: GPA (Granulomatosis with Polyangiitis), MPA (Microscopic Polyangiitis), and EGPA (Eosinophilic Granulomatosis with Polyangiitis). Blood draws occurred every 3 months with a window of +/- 1 month for a total of 9 months. Additional samples were collected off schedule if a disease flare occurred.

##### Acute COVID-19 and Sepsis

Subjects hospitalized with acute COVID-19 and sepsis were consented and enrolled across two studies (IRB#808542; IRB#843758). Sepsis and acute COVID-19 subjects were consented and enrolled within 3 days of admission to the Hospital of the University of Pennsylvania as previously described (67). Those labeled as acute COVID-19 had a positive SARS-CoV-2 PCR test. All subjects were enrolled between March and December 2020. Some individuals had additional blood draws at later timepoints, up to 24 months post-infection (IRB# 844107). The second study enrolled hospitalized, acute COVID-19 subjects enrolled in the I-SPY COVID-19 Trial (42). Enrollment occurred within 72 hours of admission at 5 trial sites (Penn, University of Alabama Birmingham, University of California San Francisco, University of Colorado, and Wake Forest University Atrium Health) with confirmed SARS-CoV-2 PCR or antigen testing and requiring greater than 6 L per minute oxygen flow. Patients or their legally authorized representatives consented to be randomized to receive a backbone treatment (remdesivir and dexamethasone) alone versus backbone with one of 12 investigational treatments. Details of the trial inclusion and exclusion criteria, and the non-backbone treatment arms have been published at https://clinicaltrials.gov/study/NCT04488081. Whole blood was collected at time of admission and 7 days later. Samples from subjects enrolled at the University of Pennsylvania were processed on the day of collection. Samples from subjects enrolled at the University of Alabama at Birmingham, University of Colorado, University of California at San Francisco, and Wake Forest University were shipped to the University of Pennsylvania and processed the day of arrival.

Non-hospitalized subjects with acute SARS-CoV-2 infections were consented and enrolled at the University of Pennsylvania (IRB# 851465). All subjects had samples collected between July-September 2024. Blood draws occurred day 5-10 post-symptom onset or positive test, day 11-29 post-symptom onset or positive test, and day 30-60 post-symptom onset or positive test.

##### Healthy

Healthy adults were enrolled for longitudinal monitoring of responses to primary and secondary SARS-CoV-2 mRNA vaccination beginning in December 2020 through March 2021, as previously reported (IRB# 844642) (42, 68), as well as before and after a fourth, fifth or sixth vaccination (IRB# 851465) (69). All subjects received either Pfizer (BNT162b2) or Moderna (mRNA-1273) mRNA vaccines. For the primary/secondary vaccination study, whole blood was collected in sodium heparin tubes (green top) for most individuals at four time points: baseline (Day -14-0), ∼2 weeks after primary immunization (Day 7-15), day of secondary immunization (Day -3-0), ∼1 week after secondary immunization (Day 7- 15). One additional subject was monitored between Day 84-150 after primary immunization. Participants were self-reported healthy without ongoing chronic health conditions. For the latter study monitoring responses to a fourth, fifth, or sixth dose of SARS-CoV-2 mRNA vaccine, whole blood was collected in sodium heparin tubes between July 2024 and January 2025, at pre-immunization (Day - 14-0), ∼ 1 week post- immunization (Day 5-10), two-four weeks post-immunization (Day 11-29) and ∼6 weeks after immunization (Day 30-60) timepoints. Additional whole blood was collected at serial monitoring visits that occurred at least three months apart from a prior study visit.

Additional healthy adults were enrolled as control subjects for other SARS-CoV2-related studies. For IRB# 852269, healthy adults who reported no persistent fatigue or cognitive symptoms, and were not hospitalized in the prior six months, were recruited from University of Pennsylvania employee volunteers at the Rehabilitation Hospital and Hospital of the University of Pennsylvania Main Hospital in October 2024. For other studies (IRB# 834263 and IRB# 845061), individuals who were self-described as healthy, including a subset of those who had recovered from a non-hospitalized COVID- 19 infection, were consented and enrolled at the University of Pennsylvania between April 2020 and November 2024. For IRB# 813913, individuals participated in the Universal COVID-19 Biobank – a component of the Penn Medicine Biobank – as a part of a study about people who may or may not have been exposed to the SARS-CoV-2 virus as well as those who had developed COVID-19 disease. Samples were collected during January 2022.

##### Hidradenitis Suppurativa

Subjects with hidradenitis suppurativa were enrolled and consented at the University of Pennsylvania. Subjects were enrolled in the study (IRB# 853228) with an initial screening study visit. A second screening visit occurred 2-20 weeks after the initial screening visit if no changes were made to the biologic treatment regimen during that time. After starting a new biologic medication (either anti-TNF or anti-IL-17), subjects provided blood samples at 3 timepoints: Days 13-15 post-biologic start (targeting day 14), day 23-33 post-biologic start (targeting day 28), and Day 105-119 post-biologic start (targeting day 112). If no changes were made to biologics treatment after the second baseline blood draw, serial monitoring blood draws occurred every 3 months.

##### Preterm and Term Infants

Infants were enrolled and whole blood was collected as previously described (IRB# 852815) (70). Briefly, preterm infants were eligible for the study if they were born between 23 weeks and 0 days and 32 weeks and 6 days of gestation. After enrollment, blood was collected in K2-EDTA vacutainers and sent to the clinical lab for CBC. Residual unused blood was collected from the clinical lab 24-48 hours after CBC was completed. Blood was stored at room temperature in the clinical lab. Blood was recurrently obtained based on availability, with a minimum of 12 days between samples, until the infant was discharged or met 40 weeks corrected gestational age. Term infants were born greater than 37.0 weeks gestational age in the Well Baby Nursery or the Hospital of the University of Pennsylvania Intensive Care Nursery.

##### Kidney Graft Failure

An individual enrolled at the University of Pennsylvania participating in the CTOT-46 study, CAR-T Cell Therapy for Desensitization in Kidney Transplantation, https://clinicaltrials.gov/study/NCT06056102, was enrolled in June 2024 and underwent serial monitoring at the following timepoints that were included in the present analysis: pre-lymphodepletion, day 0 relative to CAR-T cell infusion, and then at Day 4, Day 7, Day 10, Day 14, Day 21, Week 4, and Week 8 post-CAR-T cell infusion.

##### Long COVID-19

Patients with symptomatic post-acute sequelae of SARS-CoV-2 infection (PASC) who were not initially hospitalized and were at least six-months following their initial infection as well as individuals who were at least one year past a hospitalization for a COVID-19 infection were enrolled from the Post-COVID Assessment and Recovery Clinic at the Rehabilitation Hospital at the University of Pennsylvania (IRB# 852269). Subjects had a single blood collection that occurred between December 2022 and February 2025. Additionally, a subset of subjects that were initially enrolled in a study examining hospitalized acute SARS-CoV2 infections (IRB# 808542) and others who were enrolled at 3-15 months post-infection and who were followed longitudinally for up to 24 months that did go on to develop Long COVID (IRB# 844107).

##### Lymphoma

Subjects with non-Hodgkin lymphoma (NHL) and chronic lymphocytic leukemia (CLL) were enrolled at the University of Pennsylvania between February and September 2021, as previously described (71). Peripheral blood was collected 0–90 days prior to first SARS-CoV-2 mRNA vaccination, 1–3 weeks after the first vaccine dose, 1–5 months after the second vaccine dose, and, when applicable, 1–2 weeks after third immunization.

##### Melanoma

Subjects diagnosed with melanoma and receiving the primary two-dose SARS-CoV-2 mRNA vaccine series were consented and enrolled at the University of Pennsylvania. Subjects underwent serial monitoring at the following timepoints: the day of first vaccine dose (d0), 14 days after the first dose (d14), the day of the second vaccine dose (d21), 7 days after second vaccine dose (d28), 3 months after the first vaccine dose (∼d120), and 6-9 months after the first vaccine dose (d180-270).

##### Multiple Myeloma

Individuals participating in the Phase 2, single-arm, non-inferiority study of limited-duration teclistamab (“LimiTec”) for relapsed/refractory multiple myeloma (https://clinicaltrials.gov/study/NCT05932680) had blood collected between July 2024 and February 2025 (IRB# 853459). Peripheral blood samples from participants at the University of Pennsylvania were collected at the time of study enrollment (when stopping teclistamab, an anti-BCMA/anti-CD3 bi-specific antibody therapy), at 2 months post-enrollment, 3 months post-enrollment, 6 months post-enrollment, and if the patient needed to re-initiate therapy.

##### Multiple Sclerosis

Subjects with multiple sclerosis (MS) were enrolled between April 2021 and March 2022 (IRB# 848377). In this longitudinal study, patients with MS were treated with anti-CD20 (ocrelizumab or rituximab), glatiramer acetate (GA), interferon, or S1P receptor modulators and followed before and after receiving doses of SARS-CoV-2 mRNA vaccine. Blood was collected before the first vaccine dose, 10–12 days after the first vaccine dose, either day of or up to 2 days before the second vaccine dose, 10–12 days after the second vaccine dose and 25–30 days after the second vaccine dose, pre-third dose, 7-14 days after third dose, 4 weeks after third dose, 3 months after third dose, 6 months after third dose, 9 months after third dose, 1 year after third dose, day -28 to 0 prior to the fourth vaccine dose, and 1 month post their fourth dose. Up to 14 longitudinal samples were obtained from some MS subjects.

#### 1.2 FlowCAP-IV HIV exposure cohort

We used publicly available data from the Flow Cytometry Critical Assessment of Population Identification Methods (FlowCAP)-IV challenge (57). For this cohort peripheral blood samples were analyzed from infants classified as HIV-exposed uninfected (HEU; n=20) or unexposed (UE; n=24) controls, collected at approximately 6 months of age. Samples were treated with one of seven conditions: unstimulated, PAM (Pam3CSK4), LPS (lipopolysaccharide), PG (peptidoglycan), PIC (polyinosinic:polycytidylic acid), CPG (CpG oligodeoxynucleotides), and R848 (resiquimod). The goal of the original challenge was to identify cell populations that discriminate between HEU and UE infants. Data were obtained from FlowRepository (72) (accession FR-FCM-ZZZU).

#### 1.3 PRINCE trial cohort

Data were obtained for samples from the PRINCE trial (NCT03214250), a randomized phase 2 study evaluating nivolumab (anti-PD-1) and/or sotigalimab (anti-CD40 agonist antibody) combined with gemcitabine/nab-paclitaxel chemotherapy in patients with first-line metastatic pancreatic ductal adenocarcinoma (PDAC). A total of 105 patients were analyzed for efficacy across three treatment arms: nivolumab plus chemotherapy (n=34), sotigalimab plus chemotherapy (n=36), and sotigalimab plus nivolumab plus chemotherapy (n=35). Baseline samples were collected at cycle 1, day 1 (C1D1) prior to treatment initiation. Clinical response was categorized as partial response (PR), stable disease (SD), or progressive disease (PD) according to RECIST 1.1 criteria (73). The primary endpoint of 1- year overall survival was met for the nivolumab/chemotherapy arm (57.7%, P=0.006 compared to historical 1-year OS of 35%) but was not met for sotigalimab/chemotherapy (48.1%, P=0.062) or the triple combination (41.3%, P=0.223).

### 2. Mass cytometry

#### 2.1 Data acquisition

For all samples, 300μL of whole blood was stained using the MaxPar Direct Immune Profiling Assay (Standard BioTools, Inc, South San Francisco, CA). For infant cohorts (pre-term and full-term), if available blood volume was less that 300μL, heat-inactivated pooled human serum was added to meet 300μL (70). Samples were stained in accordance with manufacturer protocols and as previously described (74). Briefly, whole blood was added to a 5mL tube containing a pellet of lyophilized antibodies. Acute COVID-19 samples were lysed with Cal-Lyse lysing solution (Standard BioTools, Inc, South San Francisco, CA). Cells were washed, followed by fixation with 1.6% PFA. Cells were incubated at 4°C overnight prior to staining with Cell-ID Intercalator-Ir. All other cohorts underwent a similar workflow as described above. However, after incubating for 30 min in the tube of lyophilized antibodies, stained whole blood was fixed with 420 μL PROT1 buffer (Smart Tube Inc, Las Vegas, NV) and cryopreserved at -80°C. Lyse, wash, and intercalator staining were performed as above after thaw at 37°C. Stained samples were collected on a CyTOF instrument with EQ4 beads (four element calibration beads, Standard BioTools, Inc). Samples were collected across multiple CyTOF machines. Aliquots of a few large donor blood draw samples were stained with the MDIPA panel and cryopreserved. A single vial was thawed as a batch control sample for each day of acquisition and run alongside other study samples to measure instrument stability over time.

#### 2.2 Data preprocessing

For the training corpus of CyTOF datasets, .fcs files were processed in R, using static gates to remove beads, dead cells and events with outlier values for the Gaussian channels (**Supplementary Fig. 14**) (75). Doublets were removed using the Cleanet algorithm (76). For the remaining events, protein expression channels were transformed using inverse hyperbolic sine with a cofactor of 5. Following MAESTRO pre-training, low quality samples were filtered from all subsequent analyses.

For CyTOF data sets (PRINCE trial), .fcs files were gated to remove beads, debris, doublets, and dead cells using the OMIQ platform (Boston, MA). The remaining protein expression channels were transformed using inverse hyperbolic sine with a cofactor of 5.

For the FlowCAP-IV cohort, fluorescence cytometry data were preprocessed following the original challenge protocol. The marker panel consisted of 10 channels: FSC-A, SSC-A, IFN-α, CD123, MHC- II, CD14, CD11c, IL-6, IL-12, and TNF-α.

#### 2.3 Manual gating and cell population annotation

For the training set, a logistic regression model was used to annotate major cell types (Neutrophils, Eosinophils, Basophils, Myeloid cells, pDCs, B cells, Plasmablasts, NK cells, T cells) as well as debris. After initial classification, some major cell types were further subsetted using logistic regression models (Myeloid cells into Classical Monocytes, Nonclassical Monocytes and mDCs; T cells into CD4, CD8, gamma delta and DN T cells; CD4 T cells into Memory and Naïve; CD8 T cells into Memory and Naïve). Finally, a gating strategy was employed to identify downstream detailed cell populations, making use of internal negative control populations to determine marker thresholds on a file-by-file basis (Non-switched memory B cells, Switched Memory B cells, Naïve B cells, CD56bright NK cells, CD57hi NK cells, CD4 CM T cells, CD4 EM1 T cells, CD4 EM2 EM3 T cells, CD4 Tregs, activated CD4 T cells, CD56hi CD57hi CD4 T cells, CD56hi CD57lo CD4 T cells, CD56lo CD57hi CD4 T cells, Tfh cells, Th1 cells, Th17 cells, Th2 cells, CD8 CM T cells, CD8 EM1 T cells, CD8 EM2 EM3 T cells, CD8 EMRA T cells, activated CD8 T cells, CD56hi CD57hi CD8 T cells, CD56hi CD57lo CD8 T cells, CD56lo CD57hi CD8 T cells, CD8 MAIT NKT cells).

The PRINCE trial dataset underwent manual gating as described in Figure S6 in (65).

### 3. MAESTRO (77)

Let *S* be a matrix of cytometry data where rows are single cells and columns are cell features. We aim to learn a function from the space of all finite cell sets to ℝ^*D*^, mapping variable-sized inputs to fixed-dimensional representations. Standard transformers are unsuitable for unordered sets due to their reliance on positional information; we instead employ specialized attention blocks without positional encodings, ensuring permutation invariance and accommodating variable cardinality. The initial layers use permutation equivariant operations to compute attention and extract features elementwise, allowing the attention mechanism to handle arbitrary input orderings. A final pooling-by-attention step aggregates set information into a permutation invariant output. An online tokenizer teacher model provides encoded targets of the full cell set for a student model that encodes only a masked subset, thereby learning a representation of all cells from partial observation.

#### 3.1 Transformer attention blocks

##### 3.1.1 Multi-head attention block

The core component of our model is the Multi-Head Attention (*MHA*) mechanism (78), which enables the model to attend to information from multiple representation subspaces simultaneously. Given an input set *S* = {*x*_1_, *x*_2_, …, *x*_*n*_} where each *x*_*i*_ ∈ ℝ^*d*^, we first project *S* into queries *Q*, keys *K*, and values *V* using learnable weight matrices *W*_Q_, *W*_*K*_, *W*_*V*_ respectively:*Q* = *SW*_Q_, *K* = *SW*_*K*_, *V* = *SW*_*V*_. The attention mechanism for a single head is defined as:

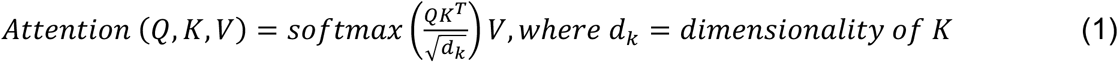

To capture diverse features, *MHA* employs *H* parallel attention heads, each computing its own attention output: *head*_ℎ_ = *Attention*(*Q*_ℎ_, *K*_ℎ_, *V*_ℎ_) for ℎ ∈ {1, …, *H*}. These outputs are concatenated and projected through a linear transformation with weight matrix *W*_0_ ∈ ℝ^*Hd*_*v*_×*d*^:

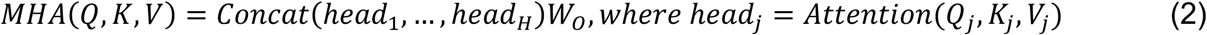

The Multi-Head Attention Block (*MAB*) integrates the *MHA* with additional components such as a cell level feed-forward neural network denoted by *FF* to enhance cell-level representations. Specifically, given set *S*_0_ and set *S*_1_, the *MAB* is defined as:

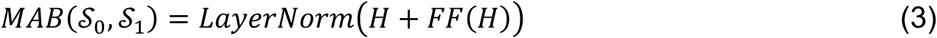

*where*,

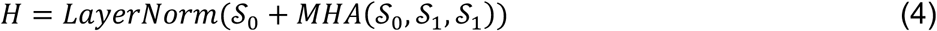

Permutation Properties of *MHA*: The *MHA* mechanism exhibits different permutation properties depending on which of its inputs are permuted, crucial for understanding how our model handles shuffled input data.

###### Theorem 1 (Permutation Properties of MHA)

Let *Q*, *K*, *V* be the query, key, and value matrices respectively, and *P_Q_*, *PKV* be permutation matrices. The Multi-Head Attention mechanism satisfies:

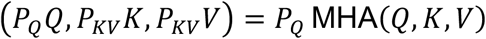

###### *Proof.* See Supplementary Information A.1

##### 3.1.2 Induced-prototype attention block

The Induced prototype attention block (*IPAB*) extends the *MAB* to efficiently handle large input sets. Based on our own studies, *IPAB*’s alone are insufficient for the number of cells in cytometry data. Given an input set **S** ∈ ℝ^*n*×*d*^ and a set of learnable inducing prototypes *I* ∈ ℝ^*m*×*d*^, where *n* is the number of elements in a set, *m* is the number of inducing prototypes, and *m* ≪ *n*, the *IPAB* is formally defined as a two-step process:

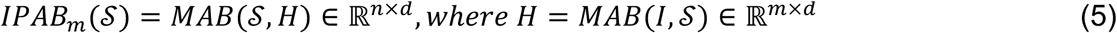

The first step transforms *I* into *H* by attending to the input set *S*. The second step uses *H*, which encodes information about *S*, to produce the final output. This structure reduces the computational complexity from *o*(*n*^2^) to *o*(*nm*), where *n* is the size of the input set. This formulation enables efficient processing of large sets assisting in the processing of large cytometry data sets.

###### Theorem 2 (Permutation Equivariance of IPAB)

The *IPAB*_*m*_(*S*) is permutation *equivariant*. Formally, for any permutation matrix *P*, it holds that:

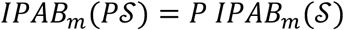

###### *Proof.* See Supplementary Information A.2

##### 3.1.3 Pooling-by-attention block

The Pooling by attention block (*PAB*) block provides a learnable aggregation mechanism for set-structured data. Given a set of input elements **S** = {*x*_1_, *x*_2_, . . ., *x*_*n*_} where each *x*_*i*_ ∈ ℝ^*d*^, and a learnable token *s* ∈ ℝ^*d*^, the *PAB* is formally defined as:

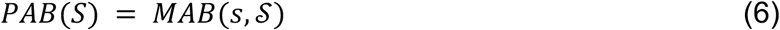

This attention-based pooling allows the model to assign varying importance to different set elements during aggregation, which is beneficial for identifying and focusing on the most relevant cell populations or features in cytometry data. This module is crucial to the permutation invariance property of MAESTRO, allowing for the collapse of hundreds of thousands of cells into a single, learnable token.

###### Theorem 3 (Permutation Invariance of PAB)

The *PAB* operation *PAB*(*S*) is permutation invariant. Formally, for any permutation *π* of the input set **S**, it holds that:

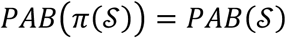

###### *Proof*. See Supplementary Information A.3

##### 3.1.4 Self-attention block

The Self-attention block (*SAB*) is a specialization of the *MAB* where the input set attends to itself. Given an input set *S* = {*x*_1_, *x*_2_, . . ., *x*_*n*_} where each *x*_*i*_ ∈ ℝ^*d*^, the *SAB* is formally defined as:

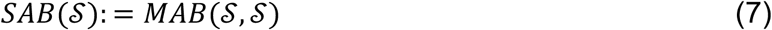

The *SAB* performs self-attention among the elements in the set, attending to itself, resulting in an output set of the same size as the input. This operation captures pairwise interactions between set elements, and stacking multiple SABs allows the model to encode higher-order interactions.

###### Theorem 4 (Permutation Equivariance of SAB)

The self-attention block *SAB*(*S*) is permutation equivariant. Formally, for any permutation matrix *P*, it holds that: SAB(PS) = P SAB(S).

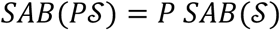

*Proof*. See Supplementary Information A.4

#### 3.2 MAESTRO Architecture

##### 3.2.1 Non-random block masking

We introduce Non-Random Masking (NRM), a technique that leverages the geometric structure of the input to construct semantically coherent masks. Given an input set *S* = {*x*_1_, . . ., *x*_*n*_} with *x*_*i*_ ∈ *R*^*d*^ and a desired mask ratio *ρ* ∈ [0, 1], NRM first centers the data and computes the first principal component via SVD, identifying the dominant axis of variation across cells. Each element is then projected onto this axis, yielding a one-dimensional ordering that reflects the primary source of phenotypic heterogeneity in the set. A contiguous block of ⌊*ρn*⌋ elements is masked from one extreme of this ordering, with the choice of which extreme randomized per sample. By removing a coherent phenotypic neighborhood rather than random individual cells, NRM challenges the model to reconstruct structured subpopulations from the remaining context, encouraging learning of global set-level relationships rather than local interpolation. As with all masking, the resulting set is shuffled to preserve permutation invariance.

##### 3.2.2 Encoder and decoder

The encoder *f* is defined as a series of nested operations, starting with a linear layer, followed by five consecutive *IPAB*s, and ending with a *PAB* operation:

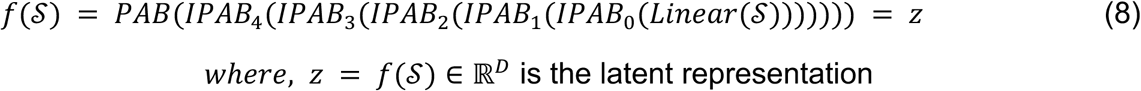

The decoder *g* takes the latent representation *z* and applies a *PAB* to unpool the vector *z* to the size of **S**, this is followed by three consecutive *SAB* operations and a linear layer to reduce the dimensionality back to the original size.

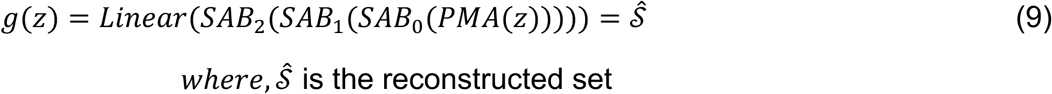

Reconstruction Loss. We employ a method called Sinkhorn Optimal Transport (SOT) for reconstruction loss (79). This approach allows us to compare the original and reconstructed sets without relying on a specific ordering of elements. We can think of this method as finding the best way to “match” elements from the original set to elements in the reconstructed set. The Sinkhorn-Knopp algorithm iteratively calculates this matching. Algorithm 2 describes the distance calculation from input to output. We define our reconstruction loss using the Sinkhorn Distance:

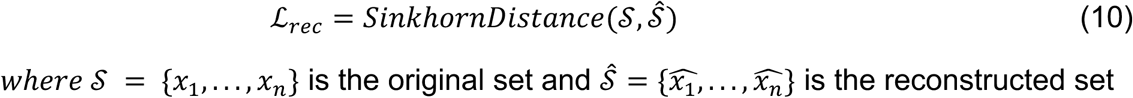

This loss function ensures that the reconstruction quality is evaluated in a permutation-invariant manner, as it only depends on how well we can match the original and reconstructed sets, not their order. For a more detailed mathematical formulation of the SOT and the reconstruction loss, refer to Supplementary Information B.

###### Algorithm 1

Sinkhorn Optimal Transport Distance

**Input**: Sets *S*, *Ŝ*, Iteration *T*

**Output**: Sinkhorn Distance *d*

1. Compute pairwise distance matrix *D_ij_* = *distance*(*x_i_* ∈ *S*, *x̂_j_* ∈ *Ŝ*)
2. Initialize 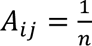 for all *i*, *j* where *A* is an *n* × *n* matrix
3. for *t* = 1 to *T* do:

a. Normalize each row of *A* such tha Σ_*i*_ *A_ij_* = 1
b. Normalize each column of *A* such tha Σ_*i*_ *A_ij_* = 1
c. Update *A_ij_* = *A_ij_* · *e^−D_ij_^*
4. Compute *d* = Σ_*ij*_ *A_ij_* · *D_ij_*

**Return: *d***

##### 3.2.3 Online tokenizer via self-distillation

MAESTRO employs a self-distillation framework with an online tokenizer to tackle the computational challenges of encoding large cytometry datasets. In this context, the teacher model acts as an online tokenizer, providing a token distribution target of the full set of cells (elements) for the student model, which operates on a masked subset. Let *S* = {*x*_1_, *x*_2_, . . ., *x*_*n*_} be the input set of cells, where each *x*_*i*_ ∈ ℝ^*d*^ represents a cell’s high-dimensional measurement. We define the student model *f*_*s*_ and the teacher model *f*_*t*_ using the encoder structure:

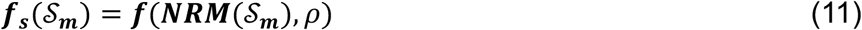

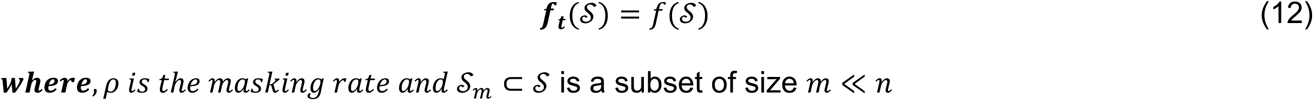

The teacher’s parameters *θ_t_* are an exponential moving average (EMA) of the student’s parameters *θ_s_*:

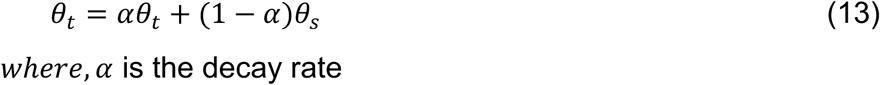

The online tokenizer (teacher) provides a stable target for the student model. The self-distillation process is achieved by minimizing the KL divergence between the softmax distributions of the latent representations produced by the student and teacher models:

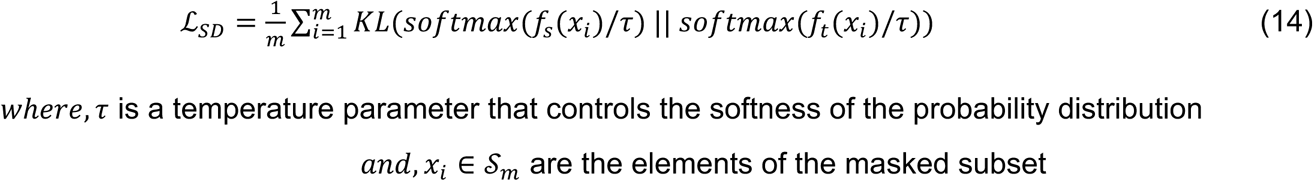

This loss drives the student’s latent representations toward those of the teacher, distilling knowledge from the complete set into a model that observes only a masked subset. Crucially, this is computationally tractable because the teacher operates on the full set without back-propagation, while the student performs gradient updates on the smaller masked subset alone.

##### 3.2.4 Overall

Using all the above components, we describe the MAESTRO algorithm here:

###### Algorithm 2

MAESTRO Model Overview

**Input**: Sets *S*, mask ratio *ρ*, EMA decay rate *α*, temperature *τ*

**Output:** Reconstructed set *Ŝ*

1. Initialize student parameters *θ_s_* and teacher parameters *θ_t_* ← *θ_s_*
2. Randomly sample subset *S_M_* ⊂ *S*
3. Apply NRM to *S_M_* to obtain masked set *S’* and mask vector *M*: (*S’*, *M*) = *NRBM*(*S_M,ρ_*)
4. Encode masked set using student encoder: *z_s_* = *f_s_*(*S’*; *θ_s_*)
5. Decode latent representation to reconstruct the set: *Ŝ* = *g*(*z_s_*; *θ_s_*)
6. Encode original set using teacher encoder: *z_t_* = *f_t_*(*S*; *θ_t_*)
7. Compute reconstruction loss: *L_rec_* = *SinkhornDistance*(*S_M_*, *Ŝ*)
8. Compute self-distillation loss: 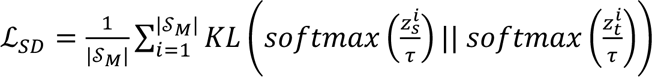
9. Compute total loss: *L* = *L_rec_* + *L_SD_*
10. Update student parameters: *θ_s_* ← *θ_s_* – *η*∇_*θ*_*s*__ *L*
11. Update teacher parameters using EMA: *θ_t_* ← *αθ_t_* + (1 – *α*)*θ_s_*

**Return:** reconstructed set *Ŝ*

#### 3.3 Training

Using 1,792 whole blood cytometry samples. We pre-train MAESTRO on four NVIDIA A100 GPUs for 100 hours at ∼12 minutes per epoch (500 epochs). Memory usage varies as each sample is of variable size, but on average the model consumes 72 GB per GPU. We train on our dataset with AdamW optimizer and a batch size of 3. The learning rate starts at 1e-4 and using a cosine annealing scheduler to a minimum of 1e-12. We use NRM to mask five different copies of our samples at masking rates of 0%, 20%, 40%, 60%, and 80%. Masking is done on the fly during pre-training. We mask by removing masked indices to the encoder, and placing masked tokens in those positions before passing through decoder. MAESTRO is configured with one attention head, a hidden size of 384, and a latent dimension of 256. Our input cells start at 30 dimensions (representing 30 different protein markers), which we use a linear layer to transform to 384 dimensions before using five IPAB (with 16 learned prototypes per IPAB) block, followed by PMA. After pooling to a single learnable seed (vector), we decode the vector by copying the pooled input to each index of the unmasked cells and use the masking token to denote indices where the original cell was masked. We follow this with a PMA block with number of seeds equal to the size of the input matrix and follow this with two SAB blocks. Finally, we use a linear layer to transform our output to its original 30 dimensions. All attention blocks use a SwiGLU activation function to learn both linear and non-linear relationships (80). Reconstruction is calculated using Sinkhorn Optimal Transport (79) with Euclidean distance for the cost matrix calculation. We align latent representations from the student and teacher model by using a non-linear projection head on the latent representation. Following the projection head we use softmax activation to convert our representations to probability distributions. We use Kullback-Leibler (KL) - Divergence to minimize the difference in distributions. The teacher model is updated with the exponential moving average of the student and a momentum value of 0.99. Student and teacher model temperatures are set to 0.1 and 0.07, respectively.

### 4. Baseline representation methods

#### 4.1 Cell type proportions

For each sample, the proportion of cells assigned to each of the 17 (or 35) gated populations was computed, yielding a fixed-dimensional composition vector. These proportion vectors served as the conventional baseline representation.

#### 4.2 MAESTRO-mini encoding baseline

To isolate the contribution of inter-cellular attention, we implemented a baseline architecture that processes cells individually through the MAESTRO encoder without the prototype-based pooling mechanism. Cell representations were aggregated via attention pooling to produce sample-level embeddings.

### 5. Embedding visualizations

Two-dimensional projections of sample embeddings were generated using Uniform Manifold Approximation and Projection (UMAP) with n_neighbors=30, min_dist=0.7, and Euclidean distance metric. For cell-level visualizations, UMAP was applied with n_neighbors=15 and min_dist varied depending on the analysis (0.0001–0.8). The UMAP implementation from the cuML library was used for GPU-accelerated computation where applicable, with random_state=253 for reproducibility.

### 6. Reconstruction evaluation

Reconstruction performance was evaluated by masking 50% of cells from held-out test samples using the cluster-based masking strategy and measuring the agreement between predicted and true marker values. For visualization (**Fig. 2a**), representative samples spanning diverse immune states were selected to illustrate reconstruction across healthy, infected (COVID-19), developmental (premature infant), malignant (lymphoma), and treatment-perturbed (multiple sclerosis) conditions. Masked cells, predicted reconstructions, and unmasked cells were jointly embedded using UMAP (n_neighbors=15, min_dist=0.2) to visualize reconstruction fidelity in a unified coordinate space.

### 7. Similarity and distance analyses

#### 7.1 Phenotype similarity

Pairwise correlations between disease phenotypes were computed by averaging MAESTRO embeddings within each phenotype and calculating correlation between phenotype centroids.

#### 7.2 Within- and between-phenotype distances

For each sample, the Pearson correlation distance (1 − Pearson r) to all other samples was computed. Distances were stratified by whether pairs shared the same phenotype (within) or differed (between).

#### 7.3 Individual stability

Euclidean distances were computed between MAESTRO embeddings of samples from the same individual across timepoints (intra-patient) and between samples from different individuals (inter-patient). Intra-patient distances were further stratified by whether the individual had healthy or disease diagnoses.

### 8. Disease classification

#### 8.1 Multi-class classification

A multi-class logistic regression classifier with L1 regularization was trained on frozen MAESTRO embeddings to discriminate among 13 disease categories. Classification was evaluated using five-fold cross-validation, with each diagnosis stratified across folds. Pre-training of MAESTRO also followed the same five-folds to assure data leakage was not contained within the model parameters themselves. Reported accuracies represent performance on held-out test samples aggregated across folds. Scikit-learn library was used (81).

#### 8.2 Few-shot classification

To evaluate sample efficiency, logistic regression classifiers were trained on progressively larger subsets of labeled data (1, 2, 5, 10, 20, … samples per class). For each training set size, multiple random subsamples were drawn, classifiers were trained, and performance was evaluated on the remaining samples. Accuracy, area under the receiver operating characteristic curve (AUROC), and area under the precision-recall curve (AUPRC) were computed for each phenotype and averaged across resampling runs.

### 9. Attention analysis

The MAESTRO pooling module assigns attention weights to each cell, quantifying the contribution to the final sample embedding. For interpretability analyses, per-cell attention weights were aggregated by summing weights within each gated population and normalizing to obtain the proportion of total attention ("attention mass") assigned to each cell type.

Attention distributions were compared across disease phenotypes (**Fig. 3c, Supplementary Fig. 7**), between response groups within treatment arms (**Fig. 5d**), and between vaccine responders and non-responders (**Fig. 4h**). Statistical comparisons used two-sided Mann-Whitney U tests with multiple testing correction by Benjamini-Hochberg false discovery rate (FDR) control. Sex-associated differences in attention (**Supplementary Fig. 8**) were evaluated using Mann-Whitney U tests with multiple testing correction by Benjamini-Hochberg false discovery rate (FDR) control. Age-associated trends (**Supplementary Fig. 9**) were assessed by Pearson correlation with FDR correction.

### 10. FlowCAP-IV HIV exposure analysis

#### 10.1 Model adaptation

MAESTRO was retrained on the FlowCAP-IV cohort to accommodate differences between fluorescence flow cytometry and mass cytometry, including the distinct 10-marker panel (FSC-A, SSC-A, IFN-α, CD123, MHC-II, CD14, CD11c, IL-6, IL-12, TNF-α). Pretraining followed the same self-supervised objectives and hyperparameters as above (**Methods 3.3**).

#### 10.2 HEU detection

Logistic regression classifiers were trained on frozen MAESTRO embeddings (256-dimensional) to discriminate HEU from UE infants. To account for the nested structure of the data (multiple stimulation conditions per individual), classification was evaluated using leave-one-individual-out cross-validation where all samples from a given individual were held out for testing while all samples from other individuals were used for training. Performance was reported as recall, precision, accuracy, F-measure, and AUROC, following the original FlowCAP-IV challenge metrics.

#### 10.3 Benchmarking

MAESTRO was compared against methods from the original FlowCAP-IV challenge as reported in (57). Results from modern cytometry deep learning approaches including CellCNN, CytoDx, and CytoSet were obtained from (82).

### 11. Vaccine response prediction

Logistic regression classifiers with L1 regularization were trained on baseline (pre-vaccination). MAESTRO embeddings to predict subsequent antibody response status. Performance was evaluated using five-fold cross-validation and compared against baseline models using: (1) demographics alone (age, sex); (2) demographics plus gated cell type proportions with logistic regression; (3) demographics plus proportions with support vector machine (SVM; RBF kernel, C=1.0); and (4) demographics plus proportions with gradient boosted trees (GradientBoostingClassifier; n_estimators=50, learning_rate=0.1, max_depth=3). Scikit-learn python library was used (81).

To assess whether predictive performance arose from compositional changes induced by vaccination, cell type proportions were compared between pre-vaccination (T1) and post-vaccination (T2) timepoints using Pearson correlation.

### 12. PRINCE trial analysis

#### 12.1 Pretraining

For PDAC response prediction, MAESTRO was pretrained on a combined corpus including the diverse phenotype cohorts (**Fig. 1**) and samples from the PRINCE trial. The 19 markers shared between the PRINCE panel and the main cohort panel were used for samples to ensure compatibility. To assess the benefit of diverse pretraining, we additionally trained a version using only PRINCE samples.

#### 12.2 Response prediction

Treatment response (PR vs SD vs PD) was predicted using logistic regression with L1 regularization (C=3.0) on frozen MAESTRO embeddings (256-dimensional). Leave-one-out cross-validation (LOOCV) was employed: for each test patient, all other samples (including longitudinal timepoints from other patients) were used for training, and the held-out patient was evaluated using only their baseline (C1D1) embedding. Baseline models included: (1) demographics alone (treatment arm, age, sex); (2) demographics plus cell type proportions with logistic regression; (3) demographics plus proportions with SVM (RBF kernel, C=1.0); and (4) demographics plus proportions with gradient boosted trees (n_estimators=50, learning_rate=0.1, max_depth=3). All baseline models used the same LOOCV framework.

#### 12.3 Attention analysis

Attention mass distributions were compared between response groups (PR, SD, PD) within each treatment arm (nivolumab/chemo, sotigalimab/chemo, combination) using two-sided Mann-Whitney U tests with Benjamini-Hochberg FDR correction. Analyses were conducted on baseline samples only and separately on all available timepoints. Attention proportions for each cell type were normalized such that the total attention mass summed to 1.0 for each sample.

#### 12.4 Survival analysis and patient stratification

##### 12.4.1 Unsupervised clustering

Baseline MAESTRO embeddings from PRINCE patients were clustered using hierarchical clustering with Ward linkage and Euclidean distance.

##### 12.4.2 Survival analysis

Overall survival (OS) and progression-free survival (PFS) were analyzed using the Kaplan-Meier method, with survival curves generated for each cluster. Differences between clusters were assessed using the log-rank test.

##### 12.4.3 Temporal dynamics

For patients with longitudinal samples, the mean Euclidean distance ("spread") between timepoint embeddings was computed as a measure of immune trajectory magnitude. Spread was compared across survival clusters and treatment arms using one-way ANOVA.

#### 12.5 Treatment selection simulation

To evaluate the potential clinical utility of immune-guided treatment selection, we simulated a counterfactual assignment strategy. For each patient, the task-specific model predicted response probabilities for each treatment arm (nivolumab/chemo, sotigalimab/chemo, combination). Patients were assigned to the therapy with the highest predicted probability of response. Patients for whom no treatment achieved a predicted response probability above 0.5 (PR) or 0.7 (SD) or for whom all predictions indicated progressive disease were classified as “no treatment recommended."

The distribution of outcomes under model-guided assignment was compared to observed outcomes under actual trial randomization. For patients assigned to “no treatment,” observed outcomes under their actual treatment were reported to assess the model’s capacity to identify patients unlikely to benefit from available therapies.

### 13 Statistical analysis

All statistical tests were two-sided unless otherwise noted. Multiple testing correction was applied using the Benjamini-Hochberg procedure to control the false discovery rate at α=0.05. Box plots throughout display the median (center line), interquartile range (box limits), and 0.025–0.975 quantiles (whiskers); outliers beyond 1.5× IQR are shown as individual points. Permutation silhouette tests were done to calculate significance of silhouette scores. We ran 10000 permutations of randomly shuffling labels and calculating silhouette score.

Statistical analyses were performed using Python 3.12 with scipy (stats module) (83) and statsmodels (multipletests) (84). Survival analyses used the lifelines package (85).

## Acknowledgements

We thank members of the Kim, Wherry, and Immune Health labs for invaluable discussions. This work was supported by NIH grants R01AI155577 (E.J.W), P01AI117950 (E.J.W), P01AI108545 (E.J.W.), U19AI082630 (E.J.W), U19AI149680 (E.J.W.), HL145754 (E.J.W.), K08AR081929 (S.A.A.), P30CA016520, Parker Institute for Cancer Immunotherapy (to E.J.W.), P30CA016520 (R.H.V.), the Colton Center for Autoimmunity at UPenn, and generous support from the Giorgi Foundation. Work in the Wherry lab is supported by the Parker Institute for Cancer Immunotherapy.

## Contributions

M.L., A.R.G., D.K., and E.J.W. conceived the study. M.L. conducted analysis. J.K., M.I., M.L.M., J.L., C.J., and A.N. provided critical comments for model development. M.L.M., S., A.P., Z.R. conducted sample preparation and M.I. conducted data preprocessing. E.M. provided support on storing and transferring data. I.K. and K.W. organized and maintained sample metadata. E.Y., C.C., J.C., and R.H.V. provided support on PDAC samples. J.W. supervised data management. J.K., J.L., Y.N provided comments on the graphical design of the figures. M.L wrote the manuscript, and J.K., J.L., A.B., A.R.G., D.K., and E.J.W. provided critical comments and valuable edits. A.R.G., D.K., and E.J.W. supervised the study.

---

**Supplementary Fig. 1:**
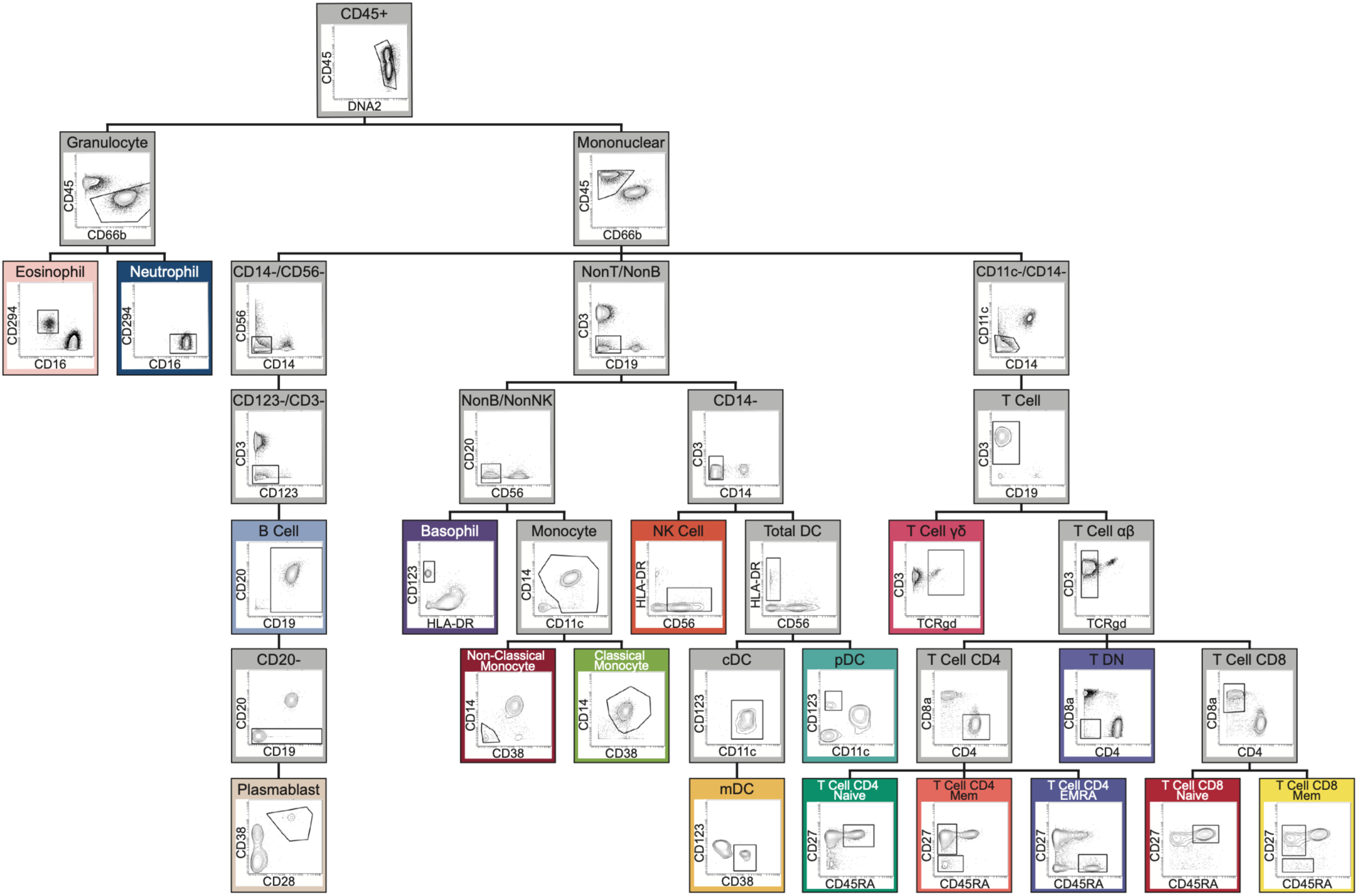
Hierarchical gating strategy for defining canonical immune populations. Sequential biaxial gating hierarchy used to annotate cytometry samples into 17 immune populations. Starting from CD45⁺ leukocytes, cells were separated into granulocyte and mononuclear compartments, then progressively gated into major lineages (T cells, B cells, NK cells, monocytes, dendritic cells, basophils, eosinophils, neutrophils) and further into subsets and differentiation states. CD4 and CD8 T cells were stratified into naive, memory, and EMRA subsets using CD27 and CD45RA; B cells were subdivided to identify plasmablasts; monocytes were separated into classical and nonclassical subsets; and dendritic cells were partitioned into cDC, pDC, and mDC. The auto-gating algorithm we use (Methods), aims to classify cell-types using this gating scheme for proportion baselines and population labels used for benchmarking and attention aggregation throughout the study.

**Supplementary Fig. 2:**
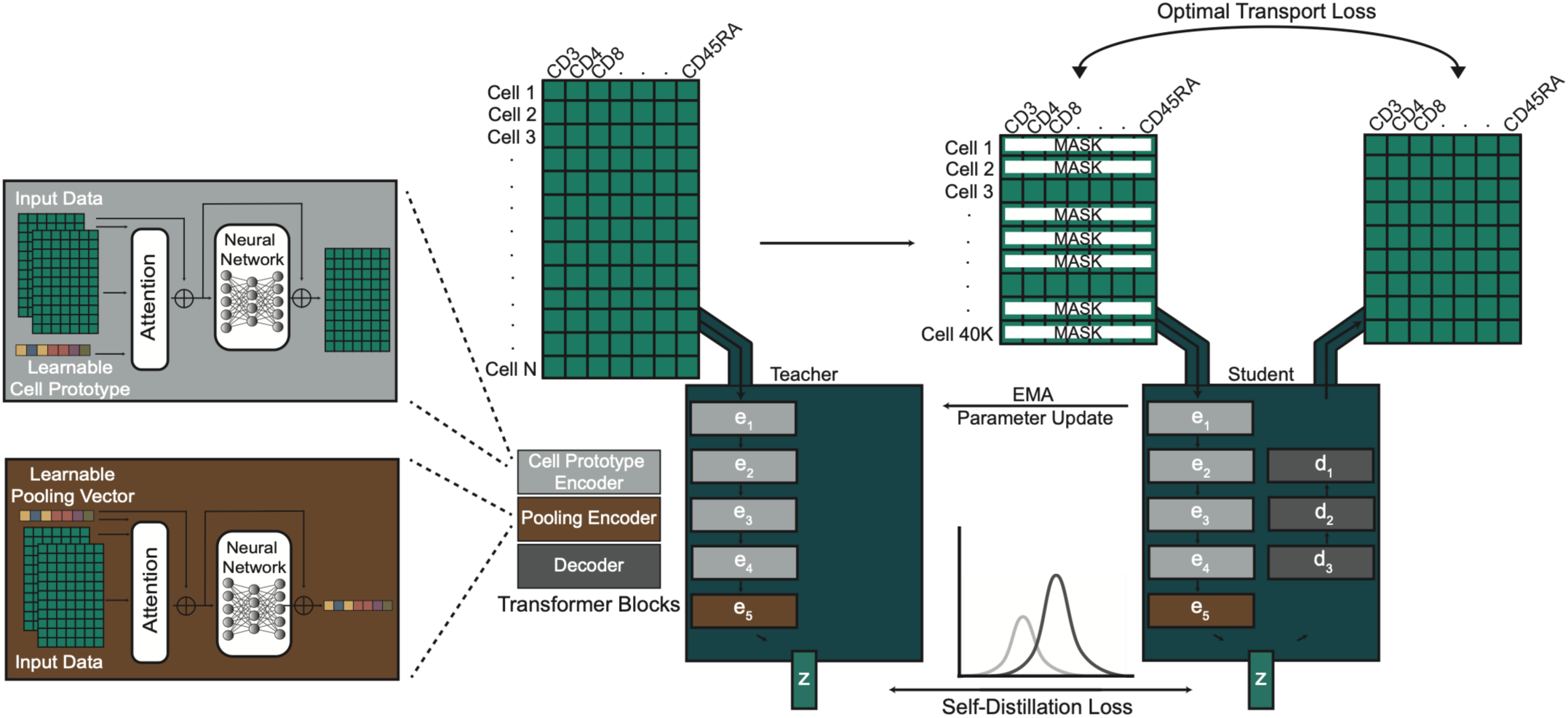
**MAESTRO architecture and self-supervised training objectives.** Schematic of the MAESTRO model showing set-based encoding of cytometry samples and the coupled self-supervised objectives used for pretraining. Each sample is represented as a cells-by-markers matrix. Cells are encoded using a prototype-based attention mechanism (Cell Prototype Encoder), in which learnable cell prototypes serve as reference points that summarize recurrent cellular patterns, enabling efficient modeling of inter-cellular relationships without all-to-all attention. Encoded representations are aggregated by a Pooling Encoder to produce a compact sample embedding z. Training uses a Student–Teacher self-distillation framework: the Teacher processes the full unmasked cell set to produce reference embeddings (e₁–e₅, z), while the Student processes masked and subsampled views and is optimized to (i) reconstruct masked cell marker values via a Decoder (d₁–d₃) using an optimal transport loss that matches predicted and true marker distributions, and (ii) align its sample embedding to the Teacher embedding via a self-distillation loss. Teacher parameters are updated as an exponential moving average (EMA) of Student parameters, providing stable training targets while enabling scaling to large cell sets.

**Supplementary Fig. 3:**
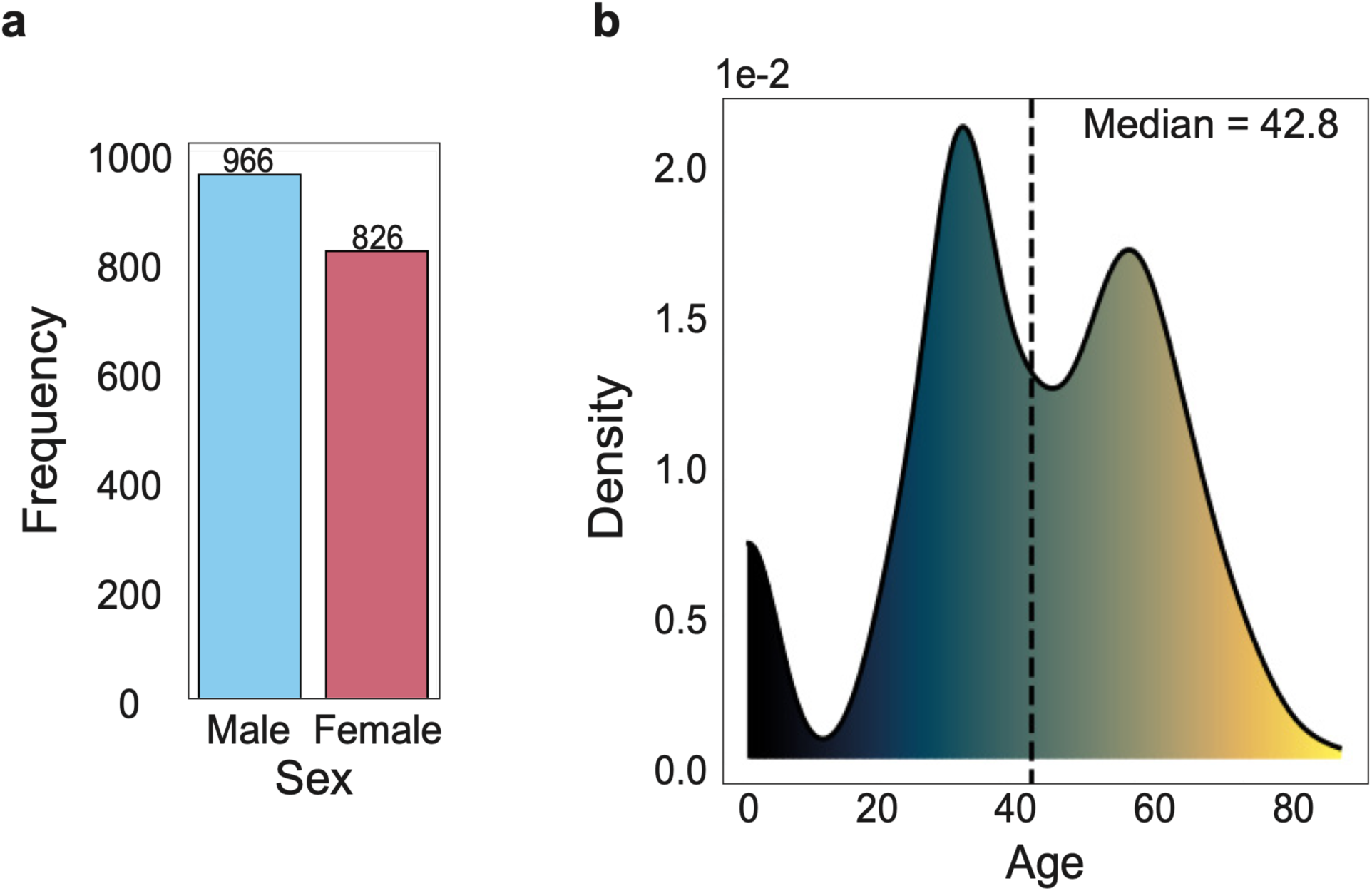
Sex and age distribution of the MAESTRO pretraining corpus. **a.** Bar plot showing the number of samples from male and female individuals in the full cytometry cohort used for self-supervised pretraining. **b.** Kernel density estimate of participant age across the full cohort, spanning premature infants to elderly adults. The dashed line indicates the cohort median age (42.8 years), and the bimodal distribution reflects the presence of both infant and adult disease cohorts.

**Supplementary Fig. 4:**
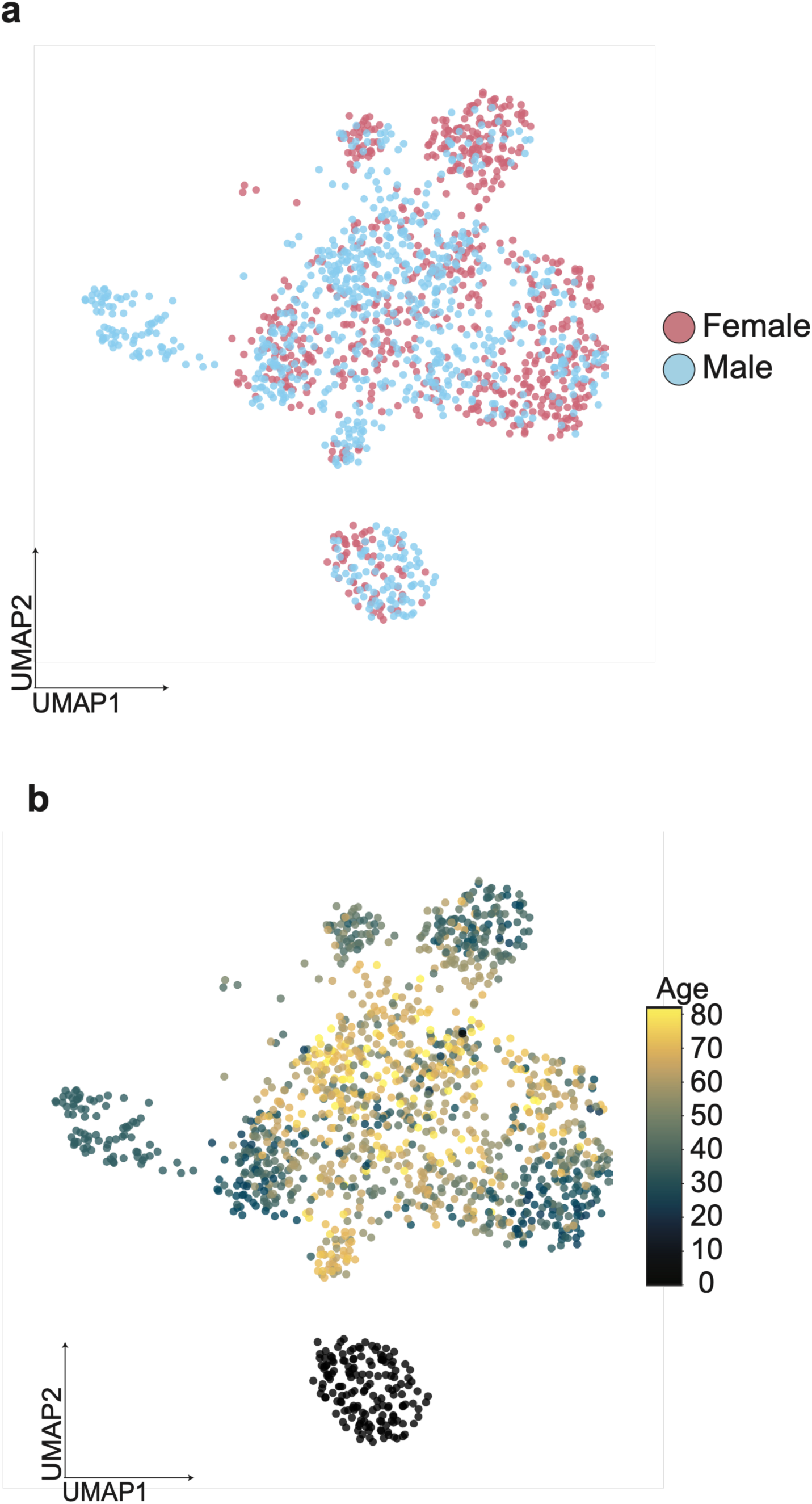
MAESTRO embeddings are structured by immune state rather than demographic covariates. **a.** UMAP projection of MAESTRO sample embeddings from **Fig. 2d** colored by sex. **b.** The same UMAP colored by age (years).

**Supplementary Fig. 5:**
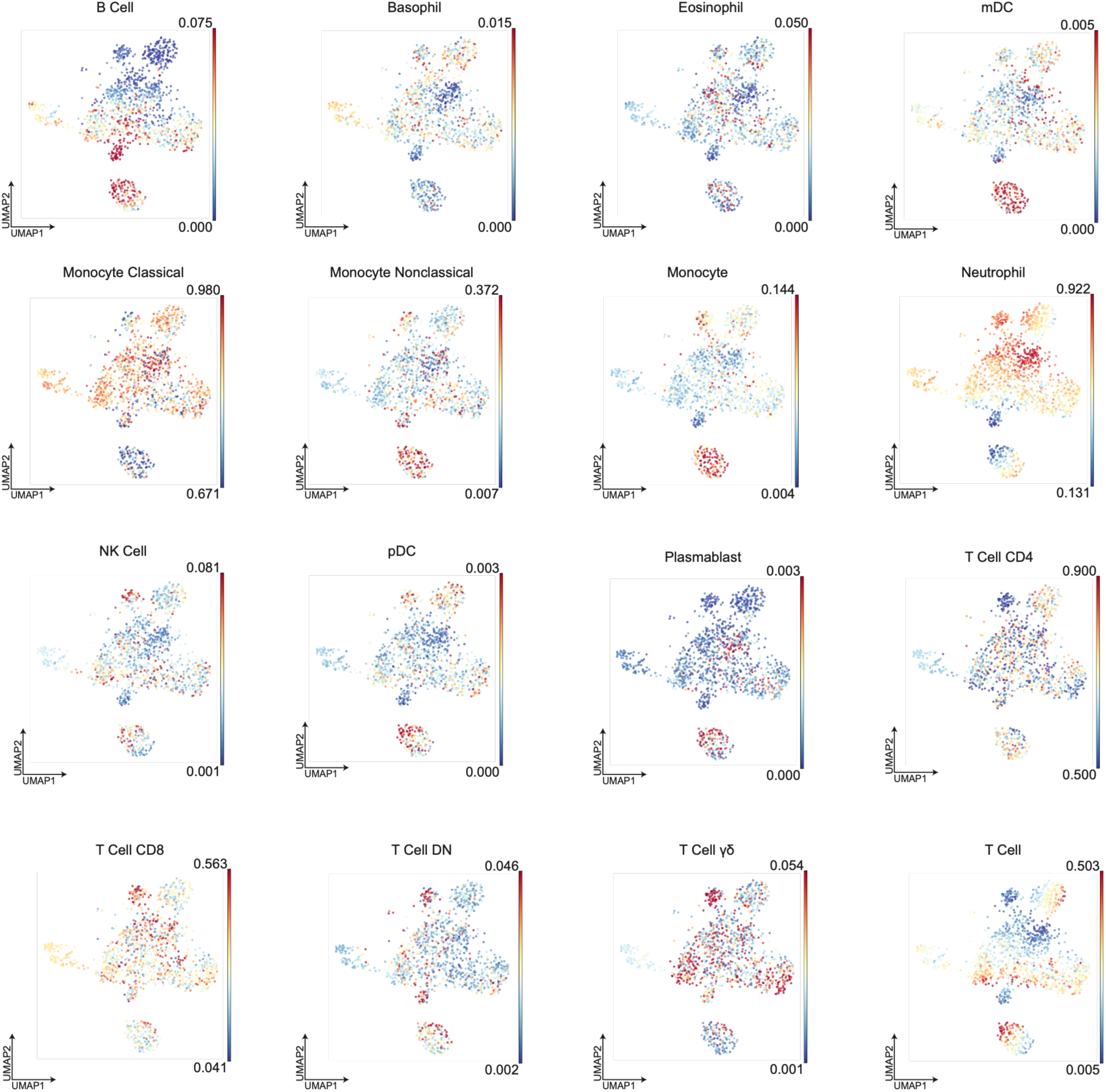
MAESTRO embedding structure is not explained by cell type proportions alone. UMAP projection of MAESTRO sample embeddings from **Fig. 2d**, colored by the proportion of each gated population within each sample (color bars indicate the range of observed proportions).

**Supplementary Fig. 6:**
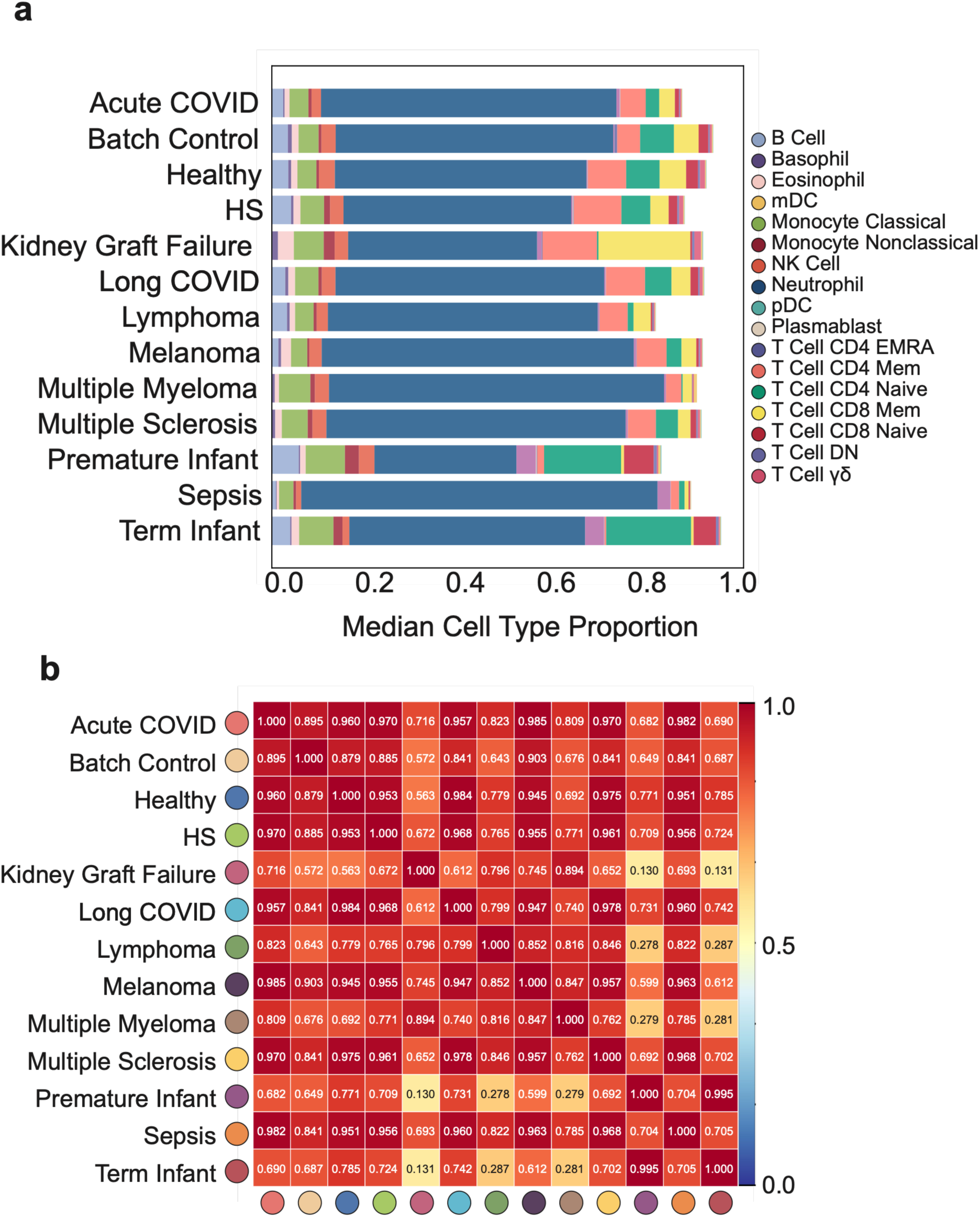
Cell type proportions remain weakly discriminative even with expanded gating resolution. **a.** Median gated cell type proportions for each disease, shown as stacked bars across the 17 canonical populations used throughout the main analyses (**Fig. 2b,e**). **b.** Pairwise cosine similarity between diseases computed from an expanded set of gated populations (35 populations, compared with 17 in **Fig. 2e**).

**Supplementary Fig. 7:**
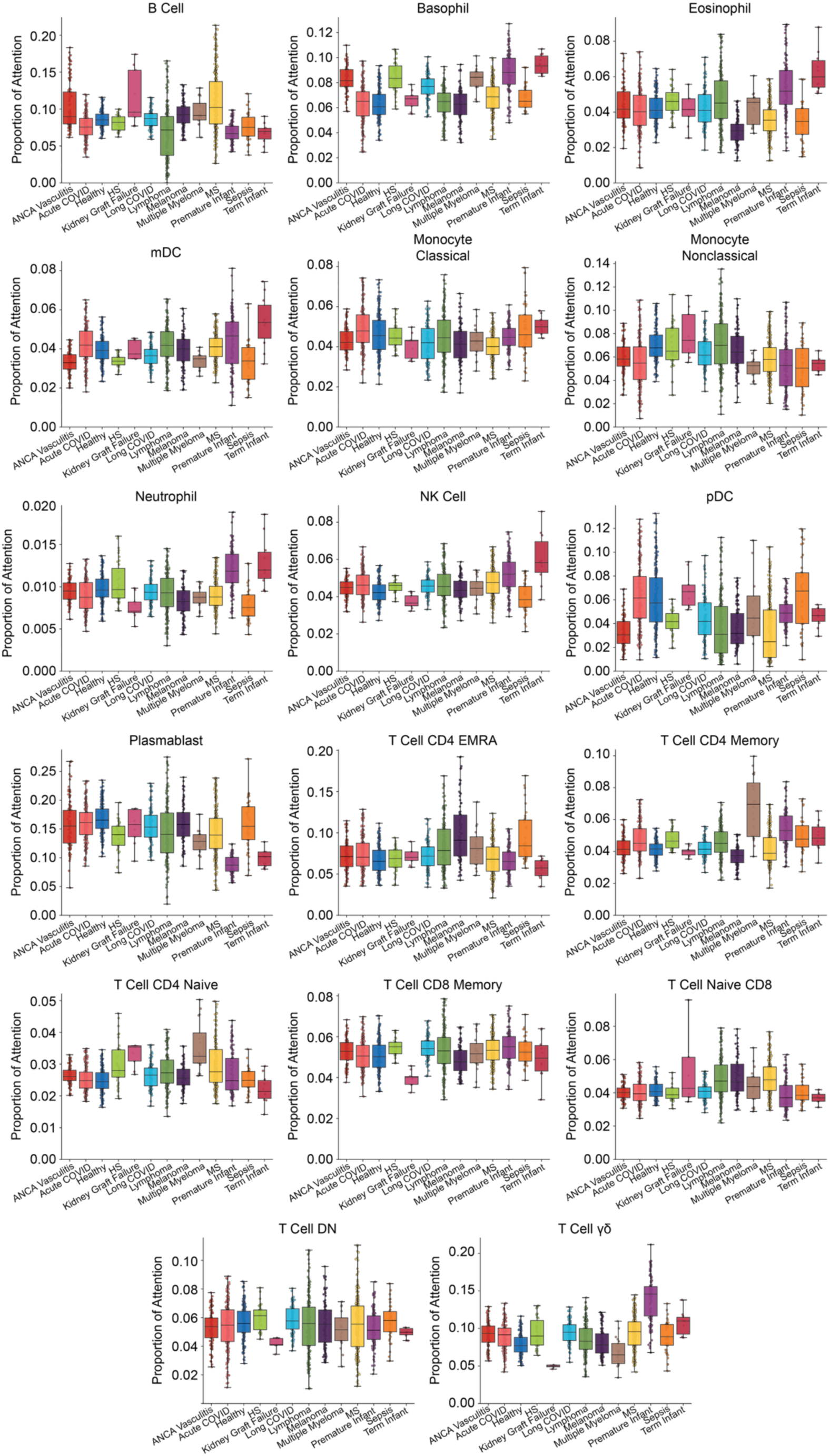
Cell-type–resolved attention allocation across disease phenotypes. Box plots show the proportion of total attention assigned to each gated population across samples from each clinical condition. For each cell type, attention proportions are summarized across samples (center line, median; box limits, first to third quartile; whiskers, 0.025 and 0.975 quantiles; points, individual samples).

**Supplementary Fig. 8:**
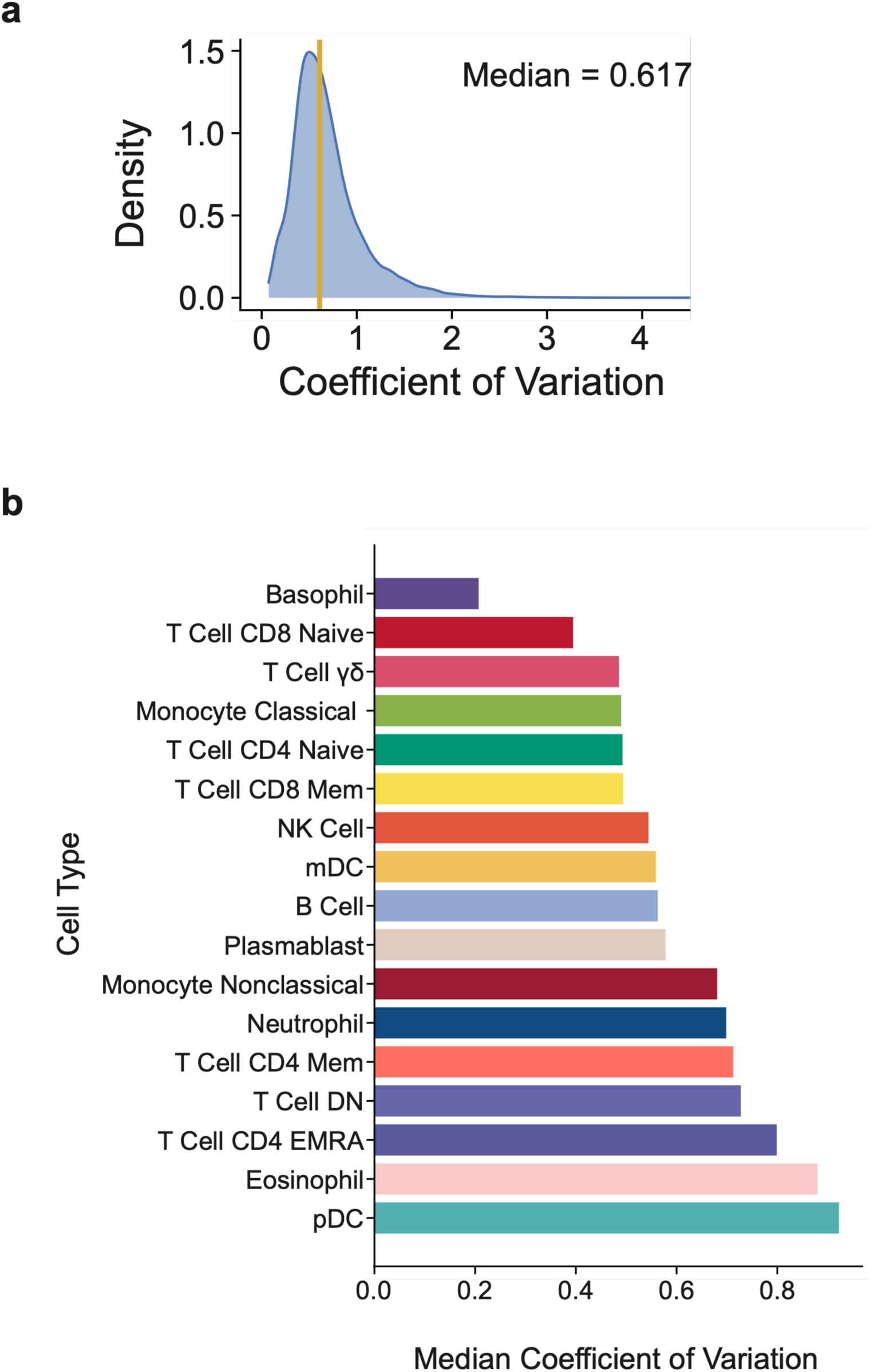
Within–cell-type variability of attention scores. **a.** Kernel density estimate of the coefficient of variation (CV) of per-cell attention weights computed within each gated population per sample, across all samples and cell types (median CV = 0.617, yellow line). **b.** Median CV per cell type across all samples, ordered by magnitude. High CV values across populations confirm that MAESTRO assigns heterogeneous attention to cells sharing the same canonical label.

**Supplementary Fig. 9:**
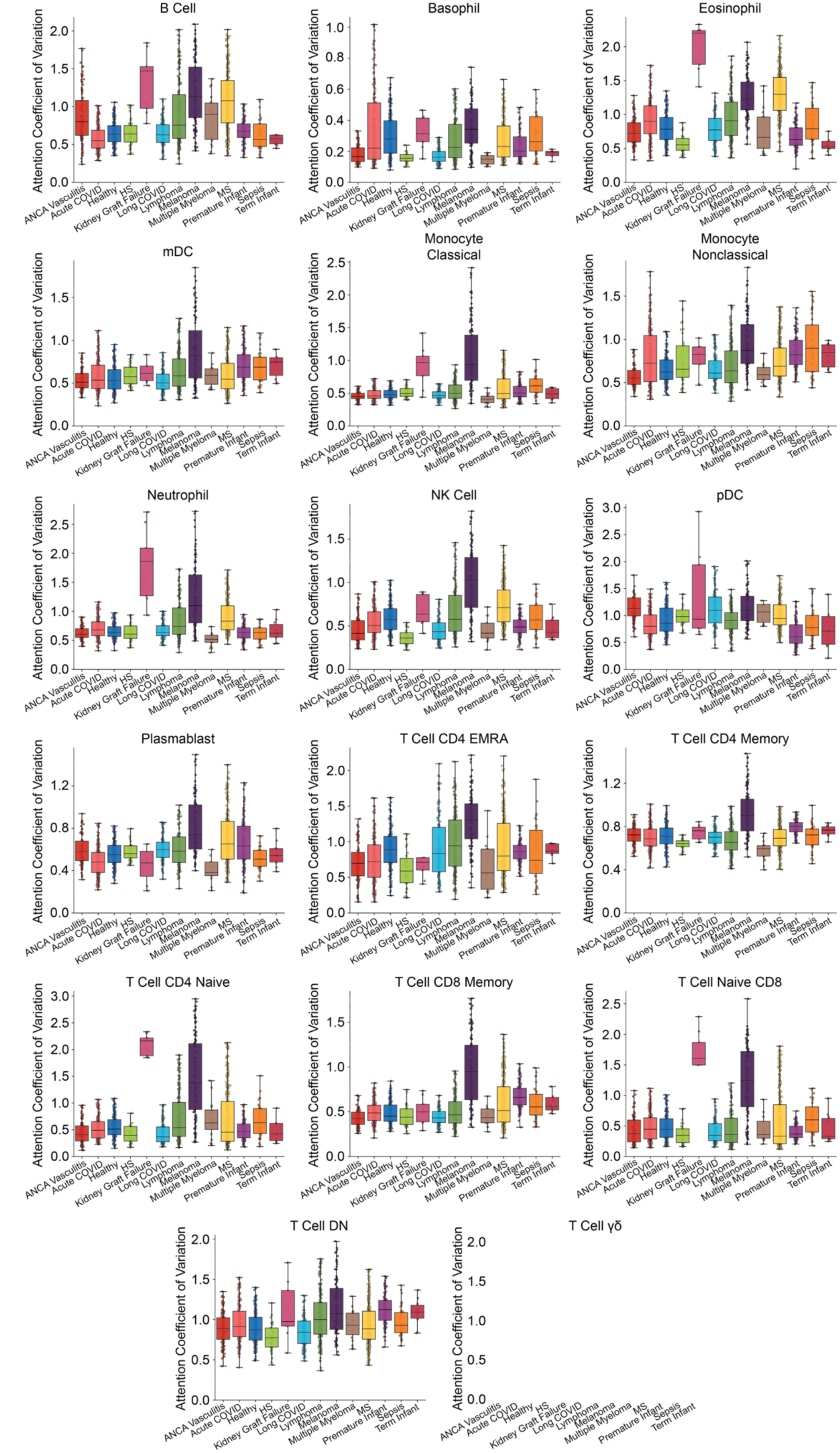
Within–cell-type attention variability across diagnostic categories. Box plots show the coefficient of variation (CV) of per-cell attention weights within each gated population, stratified by diagnosis. Each panel corresponds to one of the 17 canonical cell types; each box summarizes the distribution of CV values across samples within a diagnosis. Boxes show median (center line), interquartile range (box), and 2.5^th^–97.5^th^ percentiles (whiskers). The degree of within– cell-type attention heterogeneity varies across conditions.

**Supplementary Fig. 10:**
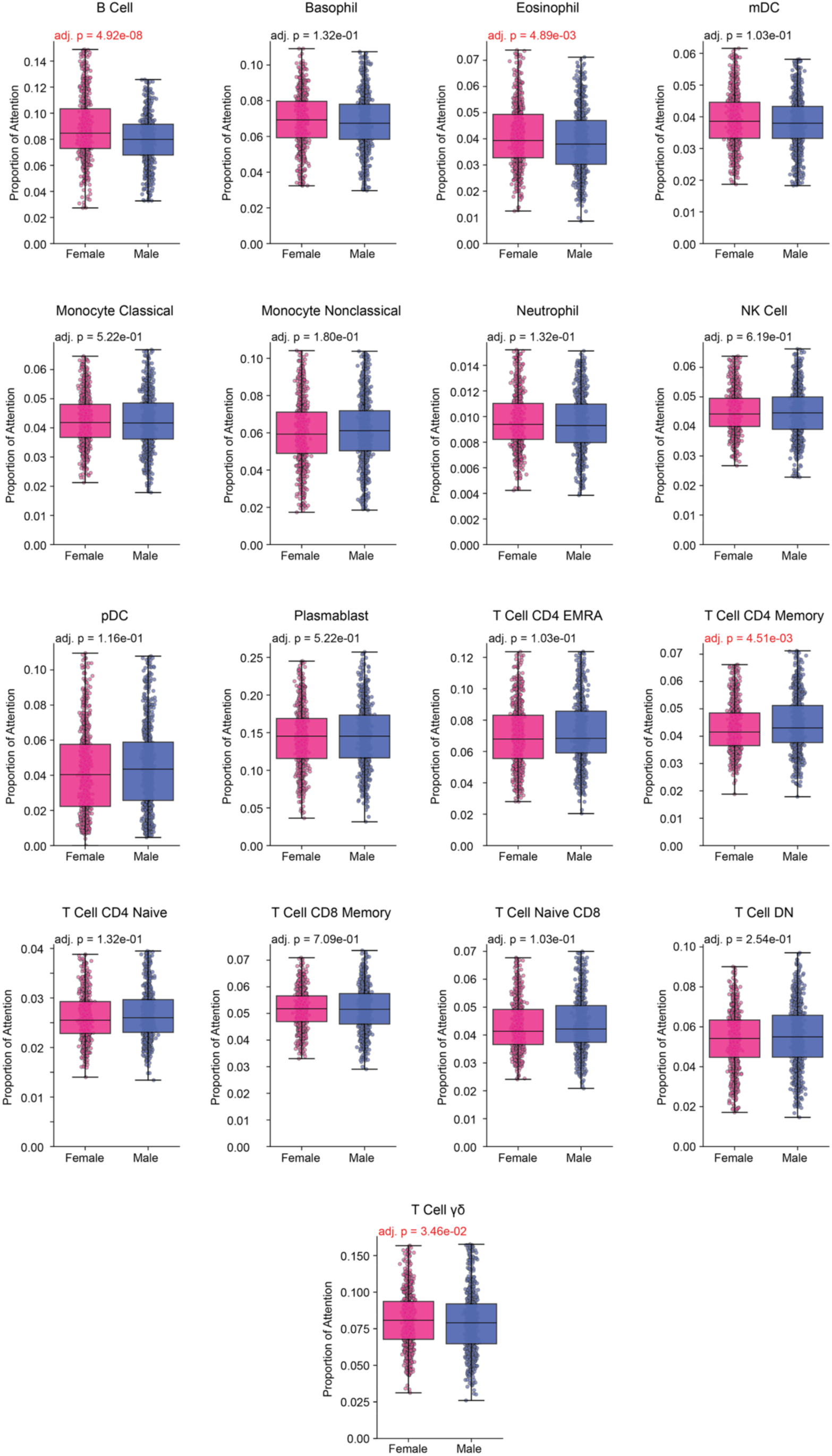
Sex-associated differences in attention allocation across immune populations. Box plots compare the proportion of total pooling attention assigned to each gated population between female and male samples. Each point represents one sample. Boxes show median (center line), interquartile range (box), and 2.5^th^–97.5^th^ percentiles (whiskers). Adjusted p-values from two-sided Mann–Whitney U tests are shown for each comparison.

**Supplementary Fig. 11:**
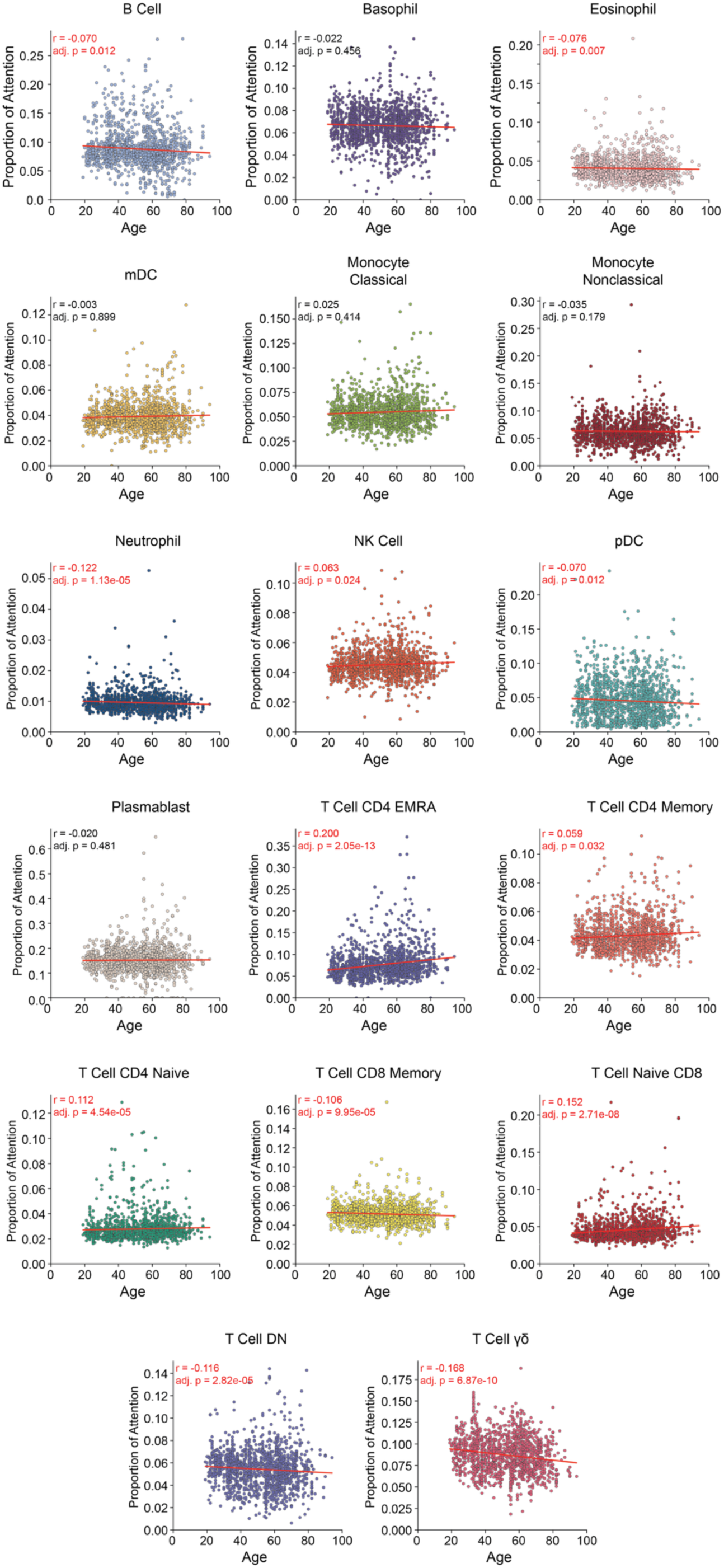
Age-associated trends in attention allocation across immune populations. Scatter plots show the proportion of total attention assigned to each gated population as a function of age, with each point representing one sample and the red line indicating a linear fit. Pearson correlation coefficients (r) and multiple-testing–adjusted P values are reported for each cell type.

**Supplementary Fig. 12:**
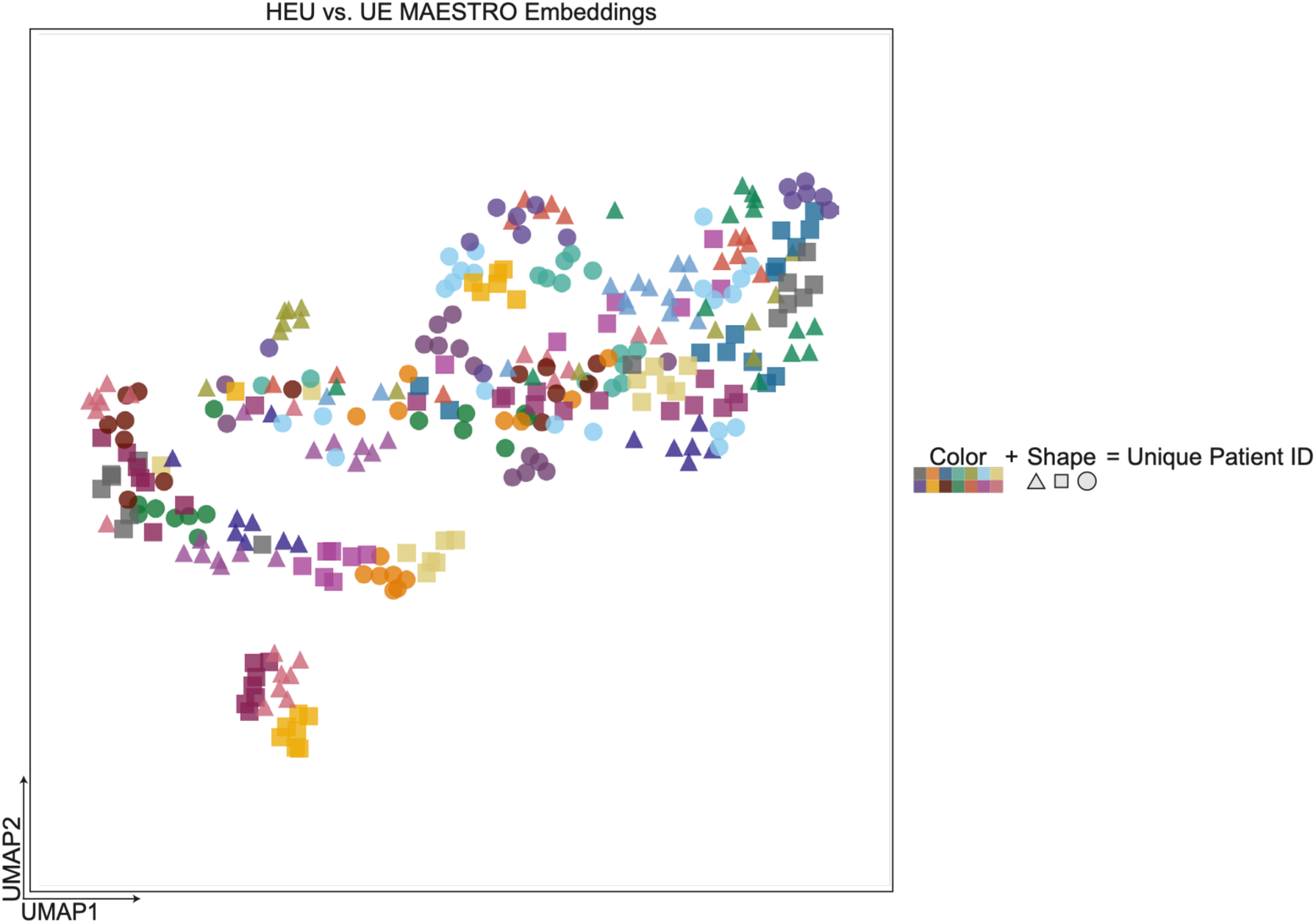
MAESTRO embeddings preserve person-specific immune fingerprints across stimulation conditions. UMAP of MAESTRO sample embeddings from the stimulation cohort. Each point represents one stimulated sample, and each unique color–shape combination denotes a single individual measured across multiple stimulation conditions. Samples from the same individual co-localize in embedding space, forming within-person clusters that are preserved across stimuli.

**Supplementary Fig. 13:**
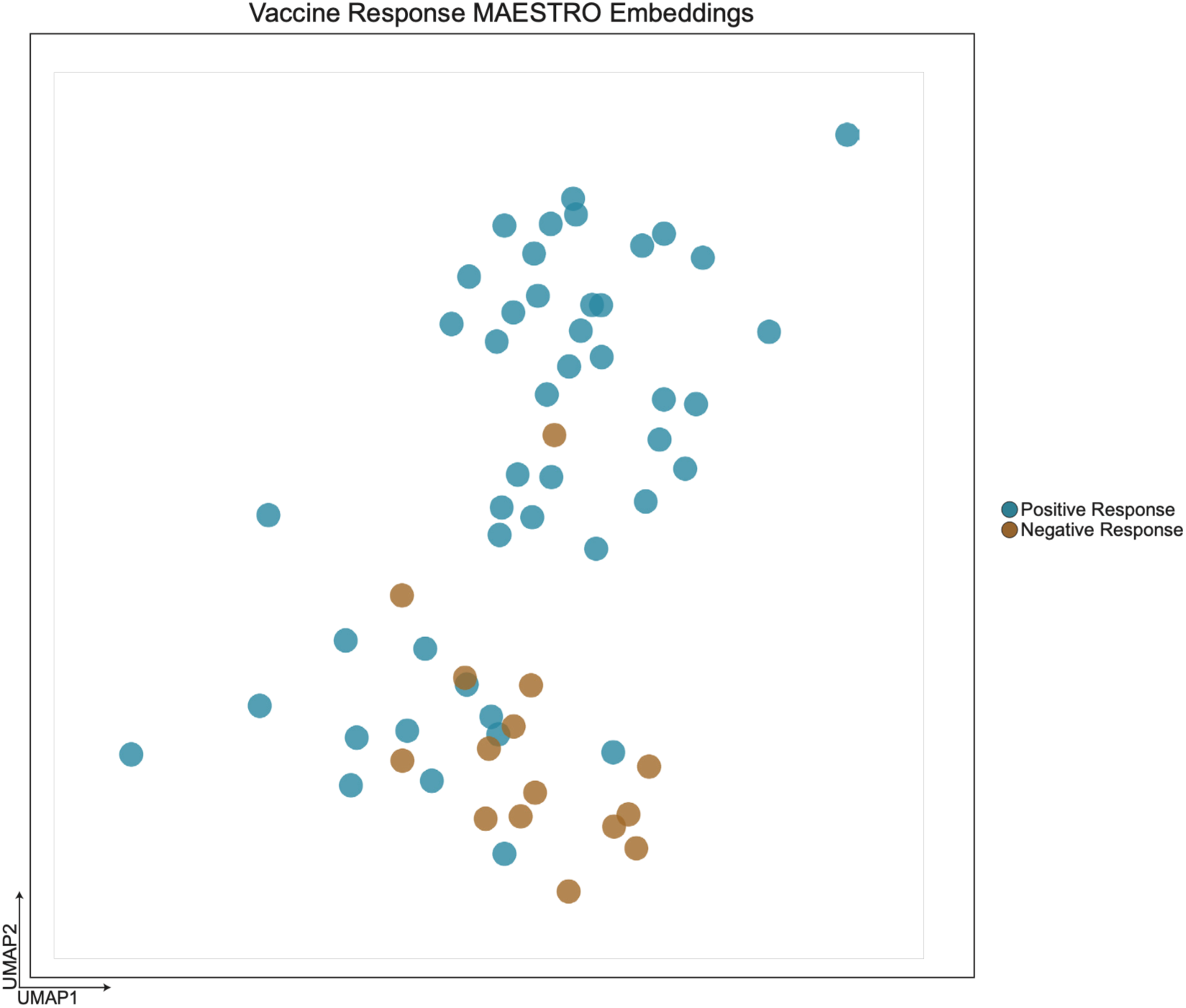
Baseline MAESTRO embeddings partially separate subsequent COVID- 19 vaccine responders. UMAP of MAESTRO sample embeddings computed from pre-vaccination (baseline) immune profiles in the COVID-19 vaccination cohort. Points represent individual baseline samples and are colored by downstream antibody response status (positive versus negative) measured at a later timepoint by IgG RBD ELISA. The embedding space shows partial segregation of responders and non-responders before vaccination, consistent with baseline immune state encoding predictive information for subsequent vaccine response.

**Supplementary Fig. 14:**
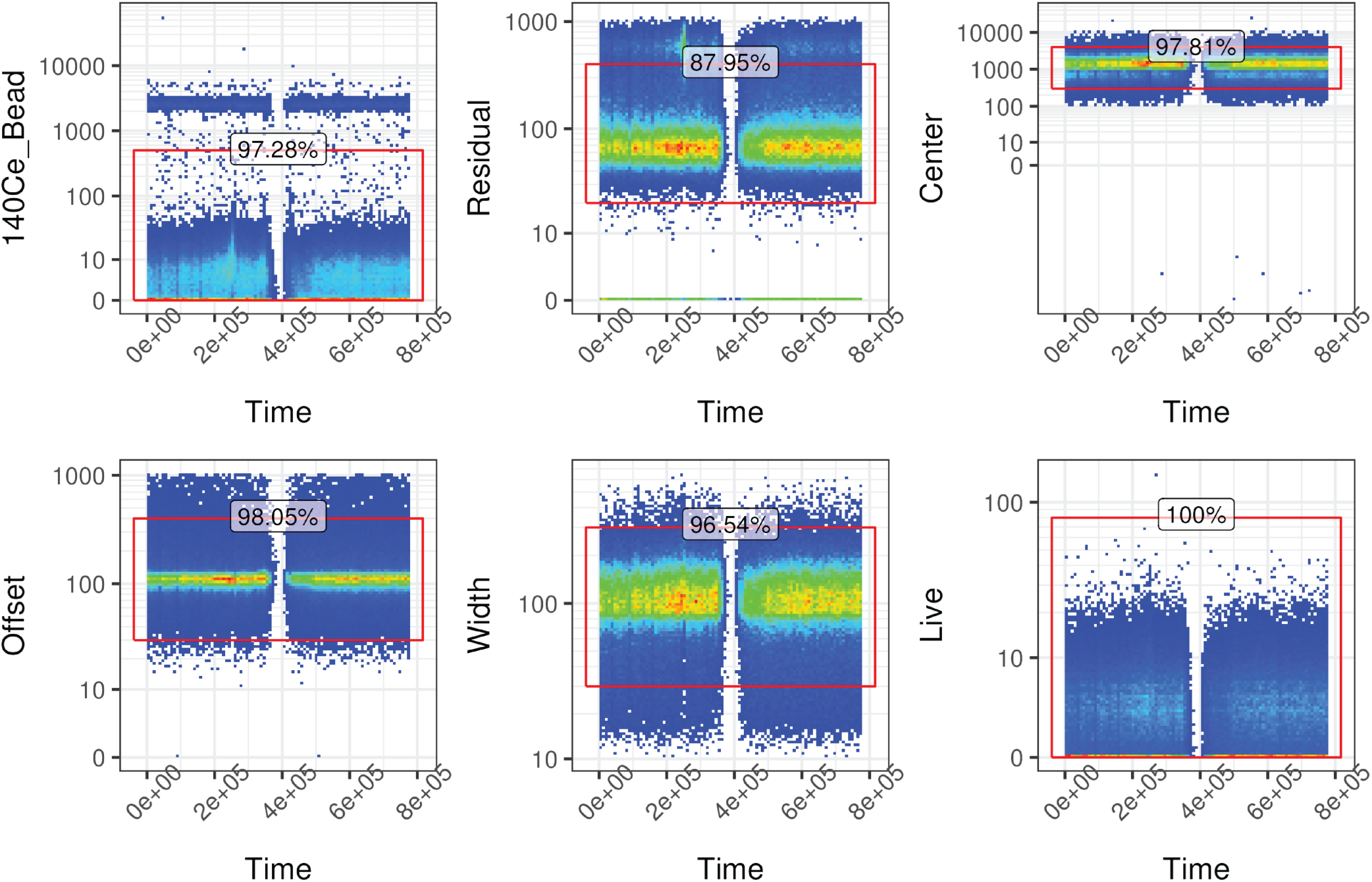
Static gating strategy for CyTOF event-level quality control. Time-resolved biaxial gates applied to raw .fcs files prior to MAESTRO pretraining to retain intact single- cell events and remove technical artifacts. Each panel plots a quality control channel against acquisition time for a representative sample; red boxes denote retained events and the percent of events passing each gate is shown above. The 140Ce bead channel was thresholded to exclude EQ4 normalization beads. The four Gaussian discrimination parameters (Residual, Center, Offset, Width) were gated to remove debris and acquisitions with anomalous signal pulse characteristics. The Live channel was gated to retain viable cells. Events passing all six gates were retained for downstream doublet removal with the Cleanet algorithm, inverse hyperbolic sine transformation with a cofactor of 5, and use as input to MAESTRO pretraining.

## Supplementary Information

### A.1 Proof of MHA Permutation Properties

We need to prove three properties of Multi-Head Attention (*MHA*):

1. Permutation Equivariance: When all inputs are permuted.
2. Permutation Equivariance for *Q*: When only the query is permuted.
3. Permutation Invariance for *K* and *V*: When only keys and values are permuted.

Let *Q* ∈ ℝ^*n_q_*×*d*^, *K*, *V* ∈ ℝ^*n*_*kv*_×*d*^ be the query, key, and value matrices respectively, and *P_q_* ∈ ℝ^*n_q_*×*n_q_*^, *P*_*qv*_ ∈ *P_kv_* ∈ ℝ^*n_kv_*×*n_kv_*^ be permutation matrices.

1. Permutation Equivariance (all inputs): When *n_q_* = *n_kv_* and *P* = *P_q_* = *P_kv_*:

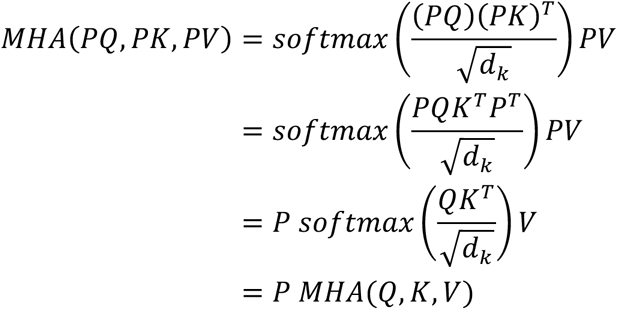

2. Permutation Equivariance for *Q*: when only *Q* is permuted:

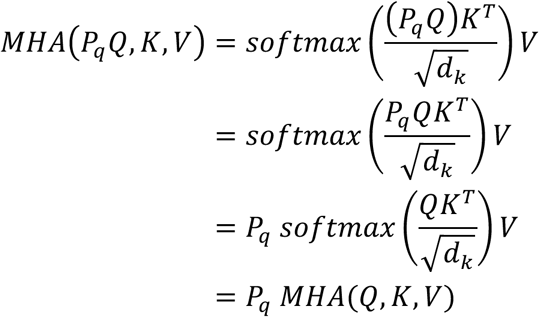

3. Permutation Invariance for *K* and *V* : when only *K* and *V* are permuted:

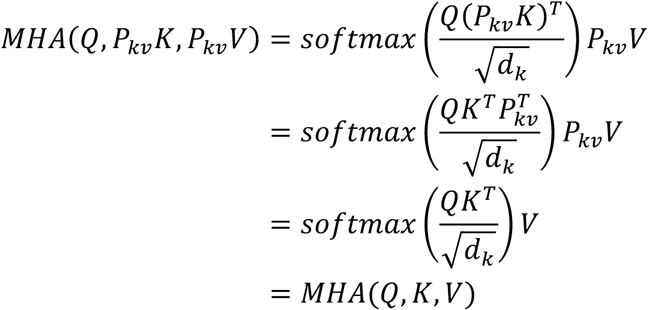

These properties together demonstrate that: a) *MHA* is permutation equivariant when all inputs are permuted equally. b) *MHA* is permutation equivariant with respect to *Q* when only *Q* is permuted. c) *MHA* is permutation invariant with respect to *K* and *V* when only *K* and *V* are permuted.

### A.2 Theorem 2 (Permutation Equivariance of IPAB)

To prove that the Induced prototype-attention block (*IPAB*) is permutation equivariant, we need to show that for any permutation matrix *P*, *IPAB*_*m*_(*PS*) = *P IPAB*_*m*_(*S*).

Let *S* ∈ ℝ^*n*×*d*^ be the input set, *I* ∈ ℝ^*m*×*d*^ be the set of learnable prototype points, and *P* ∈ ℝ^*n*×*n*^ be any permutation matrix.

Recall that *IPAB* is defined as a two-step process:

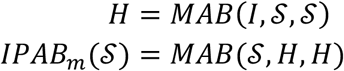

1) First, we examine how permutation affects *H*:

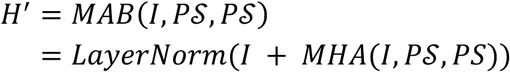

From our above *MHA* proof, we know that *MHA* is invariant when only *K* and *V* are permuted. Therefore:

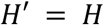

2) Now, we examine the second step:

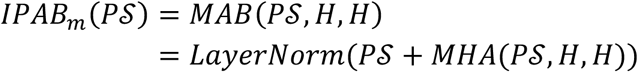

3) To show equivariance, we need to prove that this is equal to *P IPAB*_*m*_(*S*):

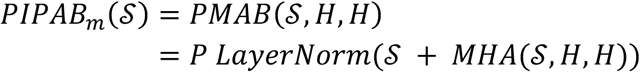

4) The equality holds because:

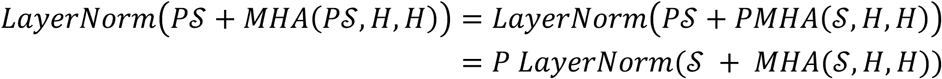

This equality holds because: a) LayerNorm is permutation equivariant b) *MHA* is permutation equivariant in its first argument (as shown in the revised *MHA* proof) c) *H* remains unchanged under permutation of **S**.

Therefore, we have shown that *IPAB*_*m*_(*PS*) = *P IPAB*_*m*_(*S*) for any permutation matrix *P*, proving that *IPAB* is permutation equivariant.

### A.3 Theorem 3 (Permutation Invariance of PAB)

To prove that the Pooling by attention (*PAB*) operation is permutation invariant, we need to show that for any permutation matrix *P* applied to the input set *S*, *PMA*(*PS*) = *PMA*(*S*).

Let *S* ∈ *R*^*n*×*d*^ be the input set, *s* ∈ *R*^1×*d*^ be the learnable token, and *P* ∈ ℝ^*n*×*n*^ be any permutation matrix. Recall that *PAB* is defined as:

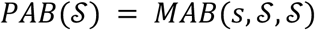

1) First, let’s consider *PAB* applied to the permuted input:

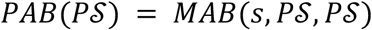

2) Expanding the *MAB* operation:

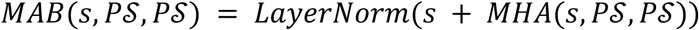

3) From our above *MHA* proof, we know that *MHA* is invariant when only *K* and *V* are permuted, which is the case here since *s* is not permuted. Therefore:

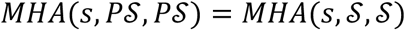

4) Consequently:

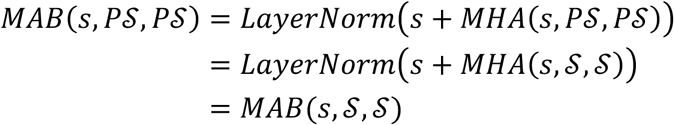

5) This shows that:

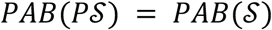

Therefore, we have shown that *PAB*(*PS*) = *PAB*(*S*) for any permutation matrix *P*, proving that *PAB* is permutation invariant.

### A.4 Theorem 4 (Permutation Equivariance of SAB)

To prove that the Self-attention block (*SAB*) is permutation equivariant, we need to show that for any permutation matrix *P*, *SAB*(*PS*) = *P SAB*(*S*).

Let *S* ∈ ℝ^*n*×*d*^ be the input set and *P* ∈ ℝ^*n*×*n*^ be any permutation matrix. Recall that *SAB* is defined as:

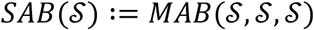

1) We start by expanding the MAB operation:

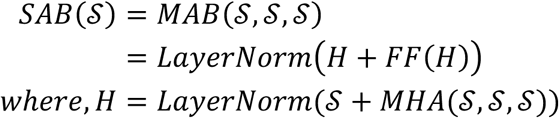

2) Now, let’s apply a permutation *P* to the input:

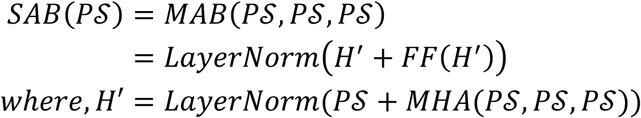

3) Using the permutation equivariance of *MHA* when all inputs are permuted (proven earlier):

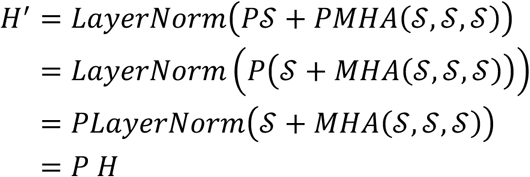

4) The feedforward network *FF* is applied elementwise, so it’s permutation equivariant:

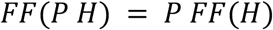

5) Combining these results

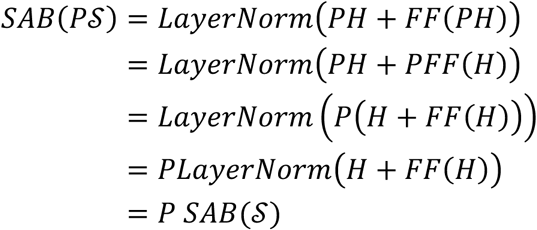

Therefore, we have shown that *SAB*(*PS*) = *P SAB*(*S*) for any permutation matrix *P*, proving that *SAB* is permutation equivariant.

#### B. Sinkhorn Distance

To ensure out reconstruction loss is permutation-invariant, we employ Sinkhorn Optimal Transport. This method allows us to compare the original and reconstructed sets without relying on a specific ordering of elements.

The key idea is to find an optimal “matching” between the original set *S* = {*x*_1_, . . ., *x*_*n*_} and the reconstructed set *Ŝ* = {*x̂*_1_, . . ., *x̂*_*n*_}. This matching is represented by a transport plan *P*.

The transport plan *P* is an *n* × *n* matrix where each element *P*_*ij*_ represents how much of an element *x*_*i*_ from the original set is “transported” to element *x̂_j_* in the reconstructed set. Formally:

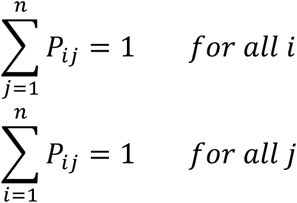

These constraints ensure that each original element is fully “distributed” across the reconstructed elements, and each reconstructed element is fully “accounted for” by the original elements.

We define a cost matrix *C*, where each element *C*_*ij*_ represents the “cost” of transporting *x*_*i*_ to *x̂_j_*:

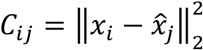

The goal is to find the optimal transport plan *P* ∗ that minimizes the total transport cost: 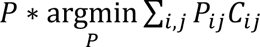 subject to the constraints on *P*

To make this optimization problem tractable, we add an entropy regularization term:

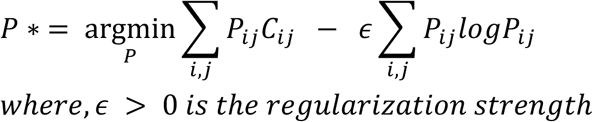

This problem can be solved efficiently using the Sinkhorn-Knopp algorithm, which iteratively updates *P* until convergence.

Finally, we define our reconstruction loss as:

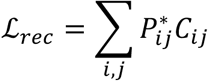

This loss function ensures that the reconstruction quality is evaluated in a permutation-invariant manner, as it only depends on the optimal matching between the original and reconstructed sets, not their order.

## References

1. Rankin LC, Artis D. Beyond Host Defense: Emerging Functions of the Immune System in Regulating Complex Tissue Physiology. Cell. 2018;173(3):554–67.

2. Mullard A. 2023 FDA approvals. Nat Rev Drug Discov. 2024;23(2):88–95.

3. Mullard A. 2024 FDA approvals. Nat Rev Drug Discov. 2025;24(2):75–82.

4. Mullard A. 2025 FDA approvals. Nat Rev Drug Discov. 2026;25(2):81–7.

5. Netea MG, Schlitzer A, Placek K, Joosten LAB, Schultze JL. Innate and Adaptive Immune Memory: an Evolutionary Continuum in the Host’s Response to Pathogens. Cell Host & Microbe. 2019;25(1):13–26.

6. Woerner J, Westbrook TM, Joo J, Shivakumar M, Venkatesh R, Cherlin T, et al. Large-scale evaluation of proteomic and polygenic risk scores reveals complementary contributions to incident disease prediction. medRxiv. 2025:2025.07.10.25331242.

7. Brodin P, Jojic V, Gao T, Bhattacharya S, Angel CJ, Furman D, et al. Variation in the human immune system is largely driven by non-heritable influences. Cell. 2015;160(1-2):37–47.

8. Cui A, Huang T, Li S, Ma A, Pérez JL, Sander C, et al. Dictionary of immune responses to cytokines at single-cell resolution. Nature. 2024;625(7994):377–84.

9. Chaplin DD. Overview of the immune response. Journal of Allergy and Clinical Immunology. 2010;125(2, Supplement 2):S3–S23.

10. Domínguez Conde C, Xu C, Jarvis LB, Rainbow DB, Wells SB, Gomes T, et al. Cross-tissue immune cell analysis reveals tissue-specific features in humans. Science. 2022;376(6594):eabl5197.

11. Dustin ML, Colman DR. Neural and Immunological Synaptic Relations. Science. 2002;2981(5594):785–9.

12. Daëron M. The immune system as a system of relations. Frontiers in Immunology. 2022;Volume 13 - 2022.

13. Altan-Bonnet G, Mukherjee R. Cytokine-mediated communication: a quantitative appraisal of immune complexity. Nature Reviews Immunology. 2019;19(4):205–17.

14. Musunuru K, Grandinette SA, Wang X, Hudson TR, Briseno K, Berry AM, et al. Patient-Specific In Vivo Gene Editing to Treat a Rare Genetic Disease. New England Journal of Medicine. 2025;392(22):2235–43.

15. Porteus MH. A New Class of Medicines through DNA Editing. New England Journal of Medicine. 2019;380(10):947–59.

16. Garraway Levi A, Lander Eric S. Lessons from the Cancer Genome. Cell. 2013;153(1):17–37.

17. Pulendran B, Davis MM. The science and medicine of human immunology. Science. 2020;369(6511):eaay4014.

18. Franceschi C, Salvioli S, Garagnani P, de Eguileor M, Monti D, Capri M. Immunobiography and the Heterogeneity of Immune Responses in the Elderly: A Focus on Inflammaging and Trained Immunity. Frontiers in Immunology. 2017;Volume 8 - 2017.

19. Song Y, Li J, Wu Y. Evolving understanding of autoimmune mechanisms and new therapeutic strategies of autoimmune disorders. Signal Transduction and Targeted Therapy. 2024;9(1):263.

20. Weiner HL. Immune mechanisms and shared immune targets in neurodegenerative diseases. Nature Reviews Neurology. 2025;21(2):67–85.

21. Hiam-Galvez KJ, Allen BM, Spitzer MH. Systemic immunity in cancer. Nature Reviews Cancer. 2021;21(6):345–59.

22. Carr EJ, Dooley J, Garcia-Perez JE, Lagou V, Lee JC, Wouters C, et al. The cellular composition of the human immune system isshaped by age and cohabitation. Nature Immunology. 2016;17(4):461– 8.

23. Brodin P, Davis MM. Human immune system variation. Nature Reviews Immunology. 2017;17(1):21–9.

24. Sparks R, Rachmaninoff N, Lau WW, Hirsch DC, Bansal N, Martins AJ, et al. A unified metric of human immune health. Nature Medicine. 2024;30(9):2461–72.

25. Davis MM. A Prescription for Human Immunology. Immunity. 2008;29(6):835–8.

26. Hao M, Gong J, Zeng X, Liu C, Guo Y, Cheng X, et al. Large-scale foundation model on single- cell transcriptomics. Nature Methods. 2024;21(8):1481–91.

27. Kalfon J, Samaran J, Peyré G, Cantini L. scPRINT: pre-training on 50 million cells allows robust gene network predictions. Nature Communications. 2025;16(1):3607.

28. Zeng Y, Xie J, Shangguan N, Wei Z, Li W, Su Y, et al. CellFM: a large-scale foundation model pre-trained on transcriptomics of 100 million human cells. Nature Communications. 2025;16(1):4679.

29. De Donno C, Hediyeh-Zadeh S, Moinfar AA, Wagenstetter M, Zappia L, Lotfollahi M, et al. Population-level integration of single-cell datasets enables multi-scale analysis across samples. Nature Methods. 2023;20(11):1683–92.

30. Tejada-Lapuerta A, Schaar AC, Gutgesell R, Palla G, Halle L, Minaeva M, et al. Nicheformer: a foundation model for single-cell and spatial omics. Nature Methods. 2025;22(12):2525–38.

31. Lopez R, Regier J, Cole MB, Jordan MI, Yosef N. Deep generative modeling for single-cell transcriptomics. Nature Methods. 2018;15(12):1053–8.

32. Rosen Y, Roohani Y, Agarwal A, Samotorčan L, Consortium TS, Quake SR, et al. Universal Cell Embeddings: A Foundation Model for Cell Biology. bioRxiv. 2023:2023.11.28.568918.

33. Heimberg G, Kuo T, DePianto DJ, Salem O, Heigl T, Diamant N, et al. A cell atlas foundation model for scalable search of similar human cells. Nature. 2025;638(8052):1085–94.

34. Yang X, Liu G, Feng G, Bu D, Wang P, Jiang J, et al. GeneCompass: deciphering universal gene regulatory mechanisms with a knowledge-informed cross-species foundation model. Cell Research. 2024;34(12):830–45.

35. Cui H, Wang C, Maan H, Pang K, Luo F, Duan N, et al. scGPT: toward building a foundation model for single-cell multi-omics using generative AI. Nature Methods. 2024;21(8):1470–80.

36. Medzhitov R. On the balance of knowledge. Nature Reviews Immunology. 2026.

37. Bendall SC, Simonds EF, Qiu P, Amir E-aD, Krutzik PO, Finck R, et al. Single-Cell Mass Cytometry of Differential Immune and Drug Responses Across a Human Hematopoietic Continuum. Science. 2011;332(6030):687–96.

38. Brixi G, Durrant MG, Ku J, Poli M, Brockman G, Chang D, et al. Genome modeling and design across all domains of life with Evo 2. bioRxiv. 2025:2025.02.18.638918.

39. Chen RJ, Ding T, Lu MY, Williamson DFK, Jaume G, Song AH, et al. Towards a general-purpose foundation model for computational pathology. Nature Medicine. 2024;30(3):850–62.

40. Hayes T, Rao R, Akin H, Sofroniew NJ, Oktay D, Lin Z, et al. Simulating 500 million years of evolution with a language model. Science. 2025;387(6736):850–8.

41. Jumper J, Evans R, Pritzel A, Green T, Figurnov M, Ronneberger O, et al. Highly accurate protein structure prediction with AlphaFold. Nature. 2021;596(7873):583–9.

42. Kim J, Ionita M, Lee M, McKeague ML, Pattekar A, Painter MM, et al. Cytometry masked autoencoder: An accurate and interpretable automated immunophenotyper. Cell Reports Medicine. 2024;5(11).

43. Bosch C, Wong JK, Paulikat M, Zapukhlyak M, Arora B, Aichmüller-Ratnaparkhe M, et al. Diversity Over Scale: Whole-Slide Image Variety Enables H&E Foundation Model Training with Fewer Patches. arXiv preprint arXiv:251110286. 2025.

44. Riaz B, Sohn S. Neutrophils in inflammatory diseases: unraveling the impact of their derived molecules and heterogeneity. Cells. 2023;12(22):2621.

45. Zhang Y, Han J. Rethinking sepsis after a two-year battle with COVID-19. Cellular & Molecular Immunology. 2022;19(11):1317–8.

46. Davis HE, McCorkell L, Vogel JM, Topol EJ. Long COVID: major findings, mechanisms and recommendations. Nature Reviews Microbiology. 2023;21(3):133–46.

47. Chen T, Kornblith S, Norouzi M, Hinton G, editors. A simple framework for contrastive learning of visual representations. International conference on machine learning; 2020: PmLR.

48. Caron M, Touvron H, Misra I, Jégou H, Mairal J, Bojanowski P, et al., editors. Emerging properties in self-supervised vision transformers. Proceedings of the IEEE/CVF international conference on computer vision; 2021.

49. Tian Y, Wang Y, Krishnan D, Tenenbaum JB, Isola P, editors. Rethinking few-shot image classification: a good embedding is all you need? European conference on computer vision; 2020: Springer.

50. Kadekar D, Agerholm R, Rizk J, Neubauer HA, Suske T, Maurer B, et al. The neonatal microenvironment programs innate gammadelta T cells through the transcription factor STAT5. J Clin Invest. 2020;130(5):2496–508.

51. Semmes EC, Chen JL, Goswami R, Burt TD, Permar SR, Fouda GG. Understanding Early-Life Adaptive Immunity to Guide Interventions for Pediatric Health. Front Immunol. 2020;11:595297.

52. Basha S, Surendran N, Pichichero M. Immune responses in neonates. Expert Rev Clin Immunol. 2014;10(9):1171–84.

53. Gibbons DL, Haque SF, Silberzahn T, Hamilton K, Langford C, Ellis P, et al. Neonates harbour highly active gammadelta T cells with selective impairments in preterm infants. Eur J Immunol. 2009;39(7):1794–806.

54. Zeiler MD, Fergus R, editors. Visualizing and understanding convolutional networks. European conference on computer vision; 2014: Springer.

55. Swiecki M, Colonna M. The multifaceted biology of plasmacytoid dendritic cells. Nature Reviews Immunology. 2015;15(8):471–85.

56. Bar-On L, Cohen H, Elia U, Cherry-Mimran L, Cohen O, Erez N. The mRNA component of LNP- mRNA vaccines triggers IFNAR-dependent immune activation which attenuates the adaptive immune response. Front Immunol. 2025;16:1670350.

57. Aghaeepour N, Finak G, Hoos H, Mosmann TR, Brinkman R, Gottardo R, et al. Critical assessment of automated flow cytometry data analysis techniques. Nature Methods. 2013;10(3):228– 38.

58. Arvaniti E, Claassen M. Sensitive detection of rare disease-associated cell subsets via representation learning. Nature Communications. 2017;8(1):14825.

59. Hu Z, Tang A, Singh J, Bhattacharya S, Butte AJ. A robust and interpretable end-to-end deep learning model for cytometry data. Proceedings of the National Academy of Sciences. 2020;117(35):21373–80.

60. Yi H, Stanley N, editors. Cytoset: Predicting clinical outcomes via set-modeling of cytometry data. Proceedings of the 12th ACM International Conference on Bioinformatics, Computational Biology, and Health Informatics; 2021.

61. Puga I, Cols M, Barra CM, He B, Cassis L, Gentile M, et al. B cell-helper neutrophils stimulate the diversification and production of immunoglobulin in the marginal zone of the spleen. Nat Immunol. 2011;13(2):170–80.

62. Nikolich-Zugich J. The twilight of immunity: emerging concepts in aging of the immune system. Nat Immunol. 2018;19(1):10–9.

63. Zhao Y, Tang J, Jiang K, Liu S-Y, Aicher A, Heeschen C. Liquid biopsy in pancreatic cancer– Current perspective and future outlook. Biochimica et Biophysica Acta (BBA)-Reviews on Cancer. 2023;1878(3):188868.

64. Liu L, Huang X, Shi F, Song J, Guo C, Yang J, et al. Combination therapy for pancreatic cancer: anti-PD-(L) 1-based strategy. Journal of experimental & clinical cancer research. 2022;41(1):56.

65. Padrón LJ, Maurer DM, O’Hara MH, O’Reilly EM, Wolff RA, Wainberg ZA, et al. Sotigalimab and/or nivolumab with chemotherapy in first-line metastatic pancreatic cancer: clinical and immunologic analyses from the randomized phase 2 PRINCE trial. Nature Medicine. 2022;28(6):1167–77.

66. Yuste R. From the neuron doctrine to neural networks. Nature Reviews Neuroscience. 2015;16(8):487–97.

67. Mathew D, Giles JR, Baxter AE, Oldridge DA, Greenplate AR, Wu JE, et al. Deep immune profiling of COVID-19 patients reveals distinct immunotypes with therapeutic implications. Science. 2020;369(6508):eabc8511.

68. Goel RR, Apostolidis SA, Painter MM, Mathew D, Pattekar A, Kuthuru O, et al. Distinct antibody and memory B cell responses in SARS-CoV-2 naïve and recovered individuals after mRNA vaccination. Science Immunology. 2021;6(58):eabi6950.

69. Painter MM, Johnston TS, Lundgreen KA, Santos JJS, Qin JS, Goel RR, et al. Prior vaccination promotes early activation of memory T cells and enhances immune responses during SARS-CoV-2 breakthrough infection. Nature Immunology. 2023;24(10):1711–24.

70. Fensterheim BA, McKeague M, Mathew D, Shwetank, Pattekar A, Nasta S, et al. Perturbation of the Preterm Human Immune System in Early Life. medRxiv. 2025.

71. Chong EA, Kumashie KG, Chong ER, Fabrizio J, Gupta A, Svoboda J, et al. Immunologic Predictors of Vaccine Responsiveness in Patients With Lymphoma and Chronic Lymphocytic Leukemia. The Journal of Infectious Diseases. 2024;230(1):15–27.

72. Spidlen J, Breuer K, Brinkman R. Preparing a Minimum Information about a Flow Cytometry Experiment (MIFlowCyt) compliant manuscript using the International Society for Advancement of Cytometry (ISAC) FCS file repository (FlowRepository.org). Curr Protoc Cytom. 2012;Chapter 10:Unit 10.8.

73. Eisenhauer EA, Therasse P, Bogaerts J, Schwartz LH, Sargent D, Ford R, et al. New response evaluation criteria in solid tumours: revised RECIST guideline (version 1.1). Eur J Cancer. 2009;45(2):228–47.

74. Geanon D, Lee B, Gonzalez-Kozlova E, Kelly G, Handler D, Upadhyaya B, et al. A streamlined whole blood CyTOF workflow defines a circulating immune cell signature of COVID-19. Cytometry Part A. 2021;99(5):446–61.

75. Iyer A, Hamers AAJ, Pillai AB. CyTOF® for the Masses. Frontiers in Immunology. 2022;Volume 13 - 2022.

76. Ionita M, McKeague ML, Painter MM, Mathew D, Pattekar A, Rezk A, et al. Cleanet: Robust Doublet Detection in Cytometry Data Based on Protein Expression Patterns. Cytometry A. 2025;107(11):716–29.

77. Lee ME, Kim J, Ionita M, Lee J, McKeague ML, Nam Y, et al. MAESTRO: Masked Encoding Set Transformer with Self-Distillation. The Thirteenth International Conference on Learning Representations2025.

78. Vaswani A, Shazeer N, Parmar N, Uszkoreit J, Jones L, Gomez AN, et al. Attention is all you need. Advances in neural information processing systems. 2017;30.

79. Cuturi M. Sinkhorn distances: Lightspeed computation of optimal transport. Advances in neural information processing systems. 2013;26.

80. Shazeer N. Glu variants improve transformer. arXiv preprint arXiv:200205202. 2020.

81. Pedregosa F, Varoquaux G, Gramfort A, Michel V, Thirion B, Grisel O, et al. Scikit-learn: Machine learning in Python. the Journal of machine Learning research. 2011;12:2825–30.

82. Yi H, Stanley N. CytoSet: predicting clinical outcomes via set-modeling of cytometry data. Proceedings of the 12th ACM International Conference on Bioinformatics, Computational Biology, and Health Informatics; Gainesville, Florida: Association for Computing Machinery; 2021. p. Article 47.

83. Virtanen P, Gommers R, Oliphant TE, Haberland M, Reddy T, Cournapeau D, et al. SciPy 1.0: fundamental algorithms for scientific computing in Python. Nature Methods. 2020;17(3):261–72.

84. Seabold S, Perktold J. Statsmodels: econometric and statistical modeling with python. SciPy. 2010;7(1):92–6.

85. Davidson-Pilon C. lifelines: survival analysis in Python. Journal of Open Source Software. 2019;4(40):1317.

